# CB1 receptor signaling modulates amygdalar plasticity during context-cocaine memory reconsolidation to promote subsequent cocaine seeking

**DOI:** 10.1101/2020.06.02.130419

**Authors:** Jessica A. Higginbotham, Rong Wang, Ben D. Richardson, Hiroko Shiina, Shi Min Tan, Mark A. Presker, David J. Rossi, Rita A. Fuchs

**Author notes:** Corresponding author: Dr. Rita A. Fuchs.

## Abstract

Contextual drug-associated memories precipitate craving and relapse in cocaine users. Such associative memories can be weakened through interference with memory reconsolidation, a process by which memories are maintained following memory retrieval-induced destabilization. We hypothesized that cocaine-memory reconsolidation requires cannabinoid type 1 receptor (**CB1R**) signaling based on the fundamental role of the endocannabinoid system in synaptic plasticity and emotional memory processing. Using an instrumental rat model of cocaine relapse, we evaluated whether systemic CB1R antagonism (**AM251**; 3 mg/kg, I.P.) during memory reconsolidation alters (a) subsequent drug context-induced cocaine-seeking behavior, as well as (b) cellular adaptations and (c) excitatory synaptic physiology in the basolateral amygdala (**BLA**). Systemic CB1R antagonism – during, but not after, cocaine-memory reconsolidation – reduced drug context-induced cocaine-seeking behavior three days, but not three weeks, later. CB1R antagonism also inhibited memory retrieval-associated increases in BLA zinc finger 268 (**zif268**) and activity regulated cytoskeletal-associated protein (**Arc**) immediate-early gene expression and changes in BLA α-amino-3-hydroxy-5-methyl-4-isoxazolepropionic acid receptor (**AMPAR**) and N-methyl-D-aspartate receptor (**NMDAR**) subunit phosphorylation that likely contribute to increased receptor membrane trafficking and synaptic plasticity during memory reconsolidation. Furthermore, CB1R antagonism increased memory reconsolidation-associated spontaneous excitatory post-synaptic current frequency in BLA principal neurons during memory reconsolidation. Together, these findings suggest that CB1R signaling modulates cellular and synaptic mechanisms in the BLA during cocaine-memory reconsolidation, thereby facilitating cocaine-memory maintenance. These findings identify the CB1R as a potential therapeutic target for relapse prevention.

**SIGNIFICANCE STATEMENT:** Drug relapse can be triggered by the retrieval of context-drug memories upon re-exposure to a drug-associated environment. Context-drug associative memories become destabilized upon retrieval and must be reconsolidated into long-term memory stores in order to persist. Hence, targeted interference with memory reconsolidation can weaken maladaptive context-drug memories and reduce the propensity for drug relapse. Our findings indicate that cannabinoid type 1 receptor (**CB1R**) signaling is critical for context-cocaine memory reconsolidation and subsequent drug context-induced reinstatement of cocaine-seeking behavior. Furthermore, cocaine-memory reconsolidation is associated with CB1R-dependent immediate-early gene expression and changes in excitatory synaptic proteins and physiology in the basolateral amygdala. Together, our findings provide initial support for CB1R as a potential therapeutic target for relapse prevention.

## INTRODUCTION

Exposure to drug-associated environmental stimuli triggers the retrieval of maladaptive drug memories that can precipitate drug craving and relapse in cocaine users (Childress et al., 1988, 1999; Crombag et al., 2008). Similar to other associative memories, cocaine-associated memories become labile upon retrieval (Nader et al., 2000). Retention of these memories requires memory reconsolidation processes that involve *de novo* protein synthesis (Nader et al., 2000; Fuchs et al., 2009; Wells et al., 2011a) and glutamatergic synaptic plasticity (Rao-Ruiz et al., 2015; Rich and Torregrossa, 2018). Consequently, targeted interference with memory reconsolidation can weaken contextual cocaine-associated memories and reduce the propensity for drug relapse (Fuchs et al., 2009; Ramirez et al., 2009; Wells et al., 2011, 2013, 2016; Arguello et al., 2014; Stringfield et al., 2017). Accordingly, it is important to investigate the cellular mechanisms of cocaine-memory reconsolidation with a focus on viable therapeutic targets.

Endocannabinoids are retrograde messengers that modulate excitatory and inhibitory synaptic plasticity (Castillo et al., 2012) and some forms of memory reconsolidation through the stimulation of presynaptic cannabinoid type 1 receptors (**CB1Rs**) (see Stern et al., 2018 for review). CB1R antagonism during memory reconsolidation impairs Pavlovian morphine- (De Carvalho et al., 2014), methamphetamine- (Yu et al., 2009), and nicotine- (Fang et al., 2011) conditioned place preference (**CPP**) memories. However, critical gaps remain in our understanding of CB1R involvement in memory reconsolidation. First, it is unclear whether CB1Rs regulate the reconsolidation of *cocaine* memories. Second, it is not known whether CB1Rs play similar roles in the reconsolidation of drug memories forged in instrumental versus Pavlovian paradigms, as extant literature indicates that Pavlovian and instrumental cocaine-associated memories are reconsolidated through partially distinct neural mechanisms (Miller and Marshall, 2005; Theberge et al., 2010; Wells et al., 2013). Finally, the cellular and synaptic physiological mechanisms by which CB1Rs modulate drug-memory reconsolidation have not been explored.

In the present study, we tested the hypothesis that CB1Rs are critically involved in contextual cocaine-memory reconsolidation in an instrumental model of drug relapse. First, we evaluated whether systemic CB1R antagonism during memory reconsolidation impairs cocaine-memory integrity as indicated by a subsequent, memory retrieval-dependent reduction in cocaine-seeking behavior. Second, we assessed the effects of memory reconsolidation and systemic CB1R antagonism on molecular adaptations and excitatory synaptic physiology in the basolateral amygdala (**BLA**), the critical site for protein synthesis-dependent memory reconsolidation in our model (Fuchs et al., 2009; Wells et al., 2011b). It has been established that auditory fear-memory reconsolidation requires memory retrieval-dependent, transient, NMDAR-dependent synaptic exchange of calcium-impermeable (**CI**, GluA2-containing) AMPARs with calcium-permeable (**CP**, GluA2-lacking) AMPARs in the lateral amygdala (Clem and Huganir, 2010; Hong et al., 2013; Lopez et al., 2015; Yu et al., 2016). While similar research has not explored the molecular mechanisms of contextual appetitive or aversive memory reconsolidation, we have shown that BLA protein kinase A (**PKA**) activation is necessary for cocaine-memory reconsolidation in the drug context-induced reinstatement model (Arguello et al., 2014). Other studies have demonstrated that PKA-mediated GluA1^S845^ phosphorylation enhances GluA1 synaptic recruitment (Clem and Huganir, 2010), whereas Src-family tyrosine kinase (**Src**)-mediated GluA2^Y876^ phosphorylation elicits GluA2 endocytosis (Hayashi and Huganir, 2004). Thus, elevated Ca^2+^ influx through NMDARs activates PKA and Src tyrosine kinases which promote changes in synaptic AMPAR-subunit composition that collectively mediate expression of long-term potentiation (**LTP;** He et al., 2009; Makino and Malinow, 2009). Moreover, Src-mediated GluN2B^Y1472^ phosphorylation is required for proper GluN2B synaptic localization, signaling, and amygdalar synaptic plasticity, raising another kinase control point for plasticity expression and/or initiation (Nakazawa et al., 2006). Accordingly, in the present study, we identified alterations in immediate-early gene (**IEGs**) expression and in glutamate receptor subunit expression and phosphorylation with a focus on GluA1^S845^, GluA2^Y876^, and GluN2B^Y1472^ phosphorylation. In an attempt to explore the synaptic physiological significance of these post-translational protein modifications, we determined changes in excitatory postsynaptic currents (**EPSCs**) in glutamatergic pyramidal principal neurons (**PNs**) of the BLA.

## MATERIALS AND METHODS

### Animals

Male Sprague-Dawley rats (*n* = 108; 275-300 g at the start of the experiment) were individually housed in a temperature- and humidity-controlled vivarium on a reversed light/dark cycle (lights on at 6:00 am). Rats were given *ad libitum* access to water and 20-25 g of standard rat chow per day. The housing and care of animals were conducted in accordance with guidelines defined in the *Guide for the Care and Use of Laboratory Animals* (National Research Council, 2011) and approved by the Washington State University Institutional Animal Care and Use Committee.

### Food Training and Surgery

To facilitate the acquisition of drug self-administration, rats were trained to press a lever (active lever) under a continuous food reinforcement schedule in standard operant conditioning chambers (Coulbourn Instruments, Holliston, MA) during a 16-h overnight food training session. Each active-lever response resulted in the unsignaled delivery of one food pellet (45 mg pellets; Bioserv, Flemington, NJ) under a continuous reinforcement schedule. Responses on a second (inactive) lever were recorded but had no programmed consequences. Contextual stimuli used for subsequent cocaine conditioning were not present during the food-training session.

For jugular catheter implantation, rats were fully anesthetized using ketamine hydrochloride and xylazine (100 mg/kg and 5 mg/kg, i.p., respectively; Dechara Veterinary Products, Overland Park, KS and Akorn, Lake Forest, IL) at least 24 h after food training. Jugular catheters were constructed in house and surgically implanted into the right jugular vein to facilitate cocaine self-administration. The catheters were maintained and periodically tested for patency as described previously (Fuchs et al., 2007). Rats received the non-steroidal anti-inflammatory analgesic, carprofen (5 g/kg per d, p.o.; ClearH2O, Westbrook, ME), from 24 h before until 48 h after surgery.

### Cocaine Self-Administration and Extinction Training

Rats were randomly assigned to one of two different environmental contexts for cocaine self-administration training. The two environmental contexts contained distinct olfactory, auditory, visual, and tactile stimuli, as previously described (Fuchs et al., 2007). Daily 2-h training sessions were conducted in one of the two environmental contexts during the rats’ dark cycle. During training sessions, active-lever responses were reinforced under a fixed ratio 1 cocaine reinforcement schedule (0.15 mg cocaine hydrochloride/0.05 mL infusion, i.v.; National Institute on Drug Abuse Drug Supply Program, Research Triangle Park, NC). Cocaine infusions were delivered over 2.25 s followed by a 20-s time-out period, during which active-lever responses had no programmed consequences. Inactive-lever responses had no programmed consequences. Training continued until the rats reached the acquisition criterion (i.e., ≥ 10 cocaine infusions obtained per session on at least 10 training days). Next, rats received daily 2-h extinction training sessions in the alternate environmental context, where lever presses had no programmed consequences. The number of extinction sessions was set at seven to hold memory age constant at the time of the experimental manipulation. Immediately after extinction session 4, the rats received an i.p. injection of saline (1 mL/kg) to acclimate them to the injection procedure.

### Experiment 1: Effects of CB1R antagonism immediately after memory retrieval on drug context-induced cocaine seeking three days later

Twenty-four h after the seventh extinction session, rats were re-exposed to the cocaine-paired context for 15 min to trigger memory retrieval and reconsolidation (**Fig. 1A**). During the session, lever presses had no programmed consequences. Cocaine reinforcement was withheld to prevent acute cocaine effects on neurotransmission and endocannabinoid mobilization independent of memory destabilization (Ortinski et al., 2012; Wang et al., 2015). Immediately after the session (i.e., during the putative time of memory reconsolidation), rats received systemic administration of the CB1R antagonist/inverse agonist, *N*- (Piperidin-1-yl)-5- (4-iodophenyl)-1- (2,4-dichlorophenyl)-4-methyl-1*H*-pyrazole-3-carboxamide (**AM251**, 3 mg/kg; Sigma Aldrich, St. Louis, MO), or vehicle (**VEH**; 8% DMSO, 5% Tween80 in saline; 1 mL/kg). AM251 at this dose is sufficient to impair contextual fear learning and memory consolidation (Arenos et al., 2006). On the next day, daily 2-h extinction training sessions resumed in the designated extinction context until the rats reached the extinction criterion (i.e., ≤ 25 active-lever presses per session on two consecutive days; mean number of days to criterion = 2.0 ± 0.0 d). Lever responses in the extinction context were assessed to detect possible off-target effects of experimental manipulations on extinction memories. Twenty-four h after the last extinction session, cocaine-seeking behavior (i.e., non-reinforced lever responses) was assessed in the cocaine-paired context. Rats were euthanized immediately after the 2-h test session, and their brains were flash frozen in isopentane and stored for analysis of BLA IEG and glutamate-receptor subunit expression using western blotting, as described below. Based on the phosphorylation kinetics of glutamate receptor subunits (Clem and Huganir, 2010; Rao-Ruiz et al., 2011), these tissue samples were prepared and analyzed for total-protein levels only.

**Figure 1.**
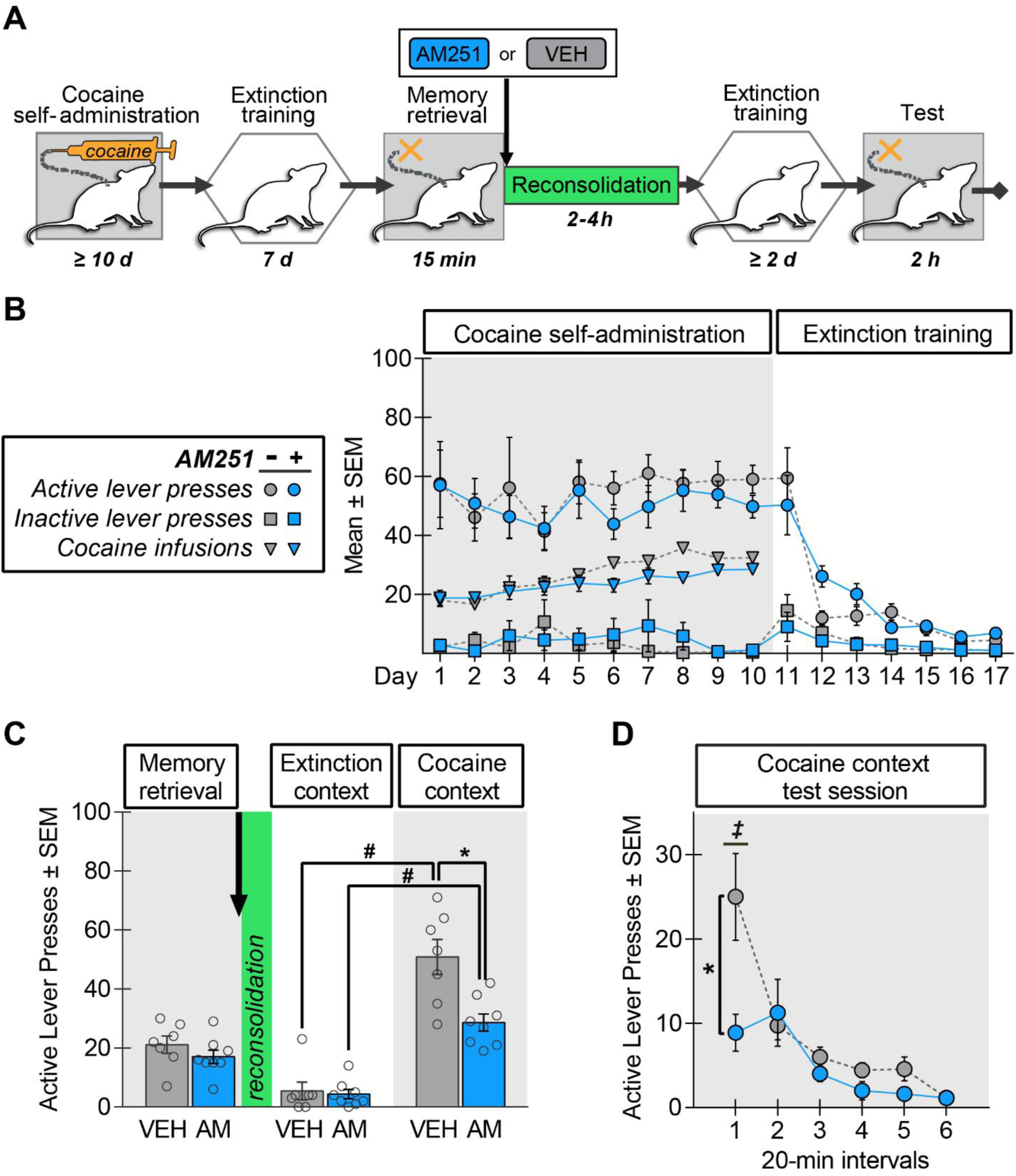
Systemic AM251 administration *during* memory reconsolidation reduces drug context-induced cocaine-seeking behavior three days later. **(A)** Experimental timeline. The 15-min memory-retrieval session was immediately followed by systemic AM251 (AM; 3 mg/kg, i.p.; *n* = 8) or VEH (*n* = 7) administration. After at least two additional daily extinction sessions, drug-seeking behavior was assessed in the cocaine-paired context. (B) Lever responses and cocaine infusions during drug self-administration (last 10 d) and extinction training. (C) Active-lever responses during the memory retrieval session in the cocaine-paired context before treatment (*arrow*) and upon first re-exposure to the extinction context and the cocaine-paired context at test. (D) Time course of active-lever presses in the cocaine-paired context at test. Symbols: ANOVA *^#^*context simple main effect, Sidak’s test, p < 0.05; *\**treatment simple main effect, Sidak’s test, p < 0.05 (Panel C), Tukey’s test, p < 0.05 (Panel D); *^‡^*time simple main effects (VEH: interval 1 > 2-6; AM251: intervals 1-2 > 4-6), Tukey’s tests, ps < 0.05.

### Experiment 2: Effects of CB1R antagonism six hours after memory retrieval on drug context-induced cocaine seeking three days later

Memory reconsolidation impairments require manipulation while the memories are labile (i.e., 2-4 h after memory retrieval; Tronson and Taylor, 2007). Experiment 2 evaluated whether any impairments in cocaine-seeking behavior in Experiment 1 reflected a memory reconsolidation deficit (**Fig. 2A**) as opposed to prolonged impairment in the expression of cocaine-seeking behavior. The procedures in Experiment 2 were identical to those in Experiment 1 except that rats received AM251 (3 mg/kg, i.p.) or VEH *six h* after the 15-min memory retrieval session in the cocaine-paired context, outside of the putative time window of memory reconsolidation. As in Experiment 1, daily 2-h extinction training sessions resumed in the extinction context after the memory retrieval session until the extinction criterion was reached (mean number of days to criterion = 2.00 ± 0.0 d). The extinction sessions were followed by a single 2-h test of cocaine-seeking behavior in the cocaine-paired context. Rats were euthanized immediately after the test session. Their brains were flash frozen in isopentane and stored for analysis of BLA IEG and glutamate receptor subunit expression using western blotting, as in Experiment 1.

**Figure 2.**
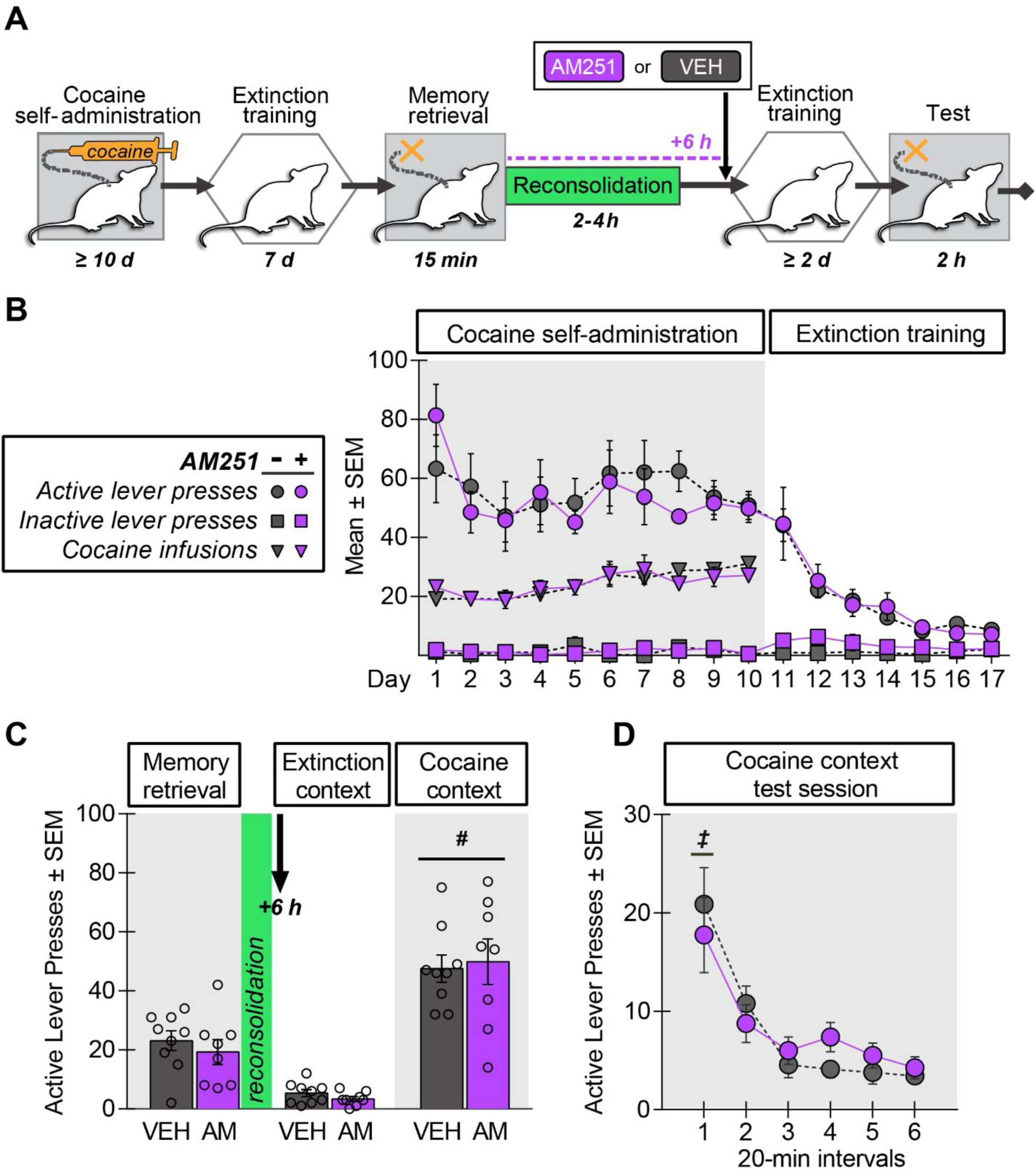
Systemic AM251 administration *after* memory reconsolidation fails to alter context-induced cocaine-seeking behavior three days later. **(A)** Experimental timeline. The 15-min memory-retrieval session was followed by systemic AM251 (AM; 3 mg/kg, i.p.; *n* = 8) or VEH (*n* = 9) administration 6 hours later (i.e., outside of the memory reconsolidation window). After at least two additional daily extinction sessions with ≤ 25 active-lever responses, drug-seeking behavior was assessed in the cocaine-paired context. **(B)** Lever responses and cocaine infusions during drug self-administration and extinction training. **(C)** Active-lever responses during the memory retrieval session in the cocaine-paired context before treatment (*arrow*) and upon first re-exposure to the extinction context and the cocaine-paired context at test. **(D)** Time course of active-lever responses across in the cocaine-paired context at test. Omnibus ANOVA effects are reported in the Results section. **Symbols**: ANOVA *^#^*context main effect, p < 0.05; *^‡^*time simple main effects (interval 1 > 2-6), Tukey’s tests, ps < 0.05.

### Experiment 3: Effects of CB1R antagonism immediately after memory retrieval on drug context-induced cocaine seeking 24 days later

Experiment 3 assessed whether any effects of AM251 on memory integrity persisted over time. The procedures were identical to those in Experiment 1 except that, after memory retrieval and pharmacological manipulation, rats remained in their home cages for 21 days (**Fig. 3A**). Daily 2-h extinction training sessions then resumed until the extinction criterion was reached (mean number of days to criterion = 2.93 ± 0.27 d), and this was followed by a 2-h test of cocaine-seeking behavior (i.e., 24 d post treatment). Rats were euthanized immediately after the test session. Their brains were flash frozen in isopentane and stored for analysis of BLA IEG and glutamate-receptor subunit expression using western blotting, as in Experiment 1.

**Figure 3.**
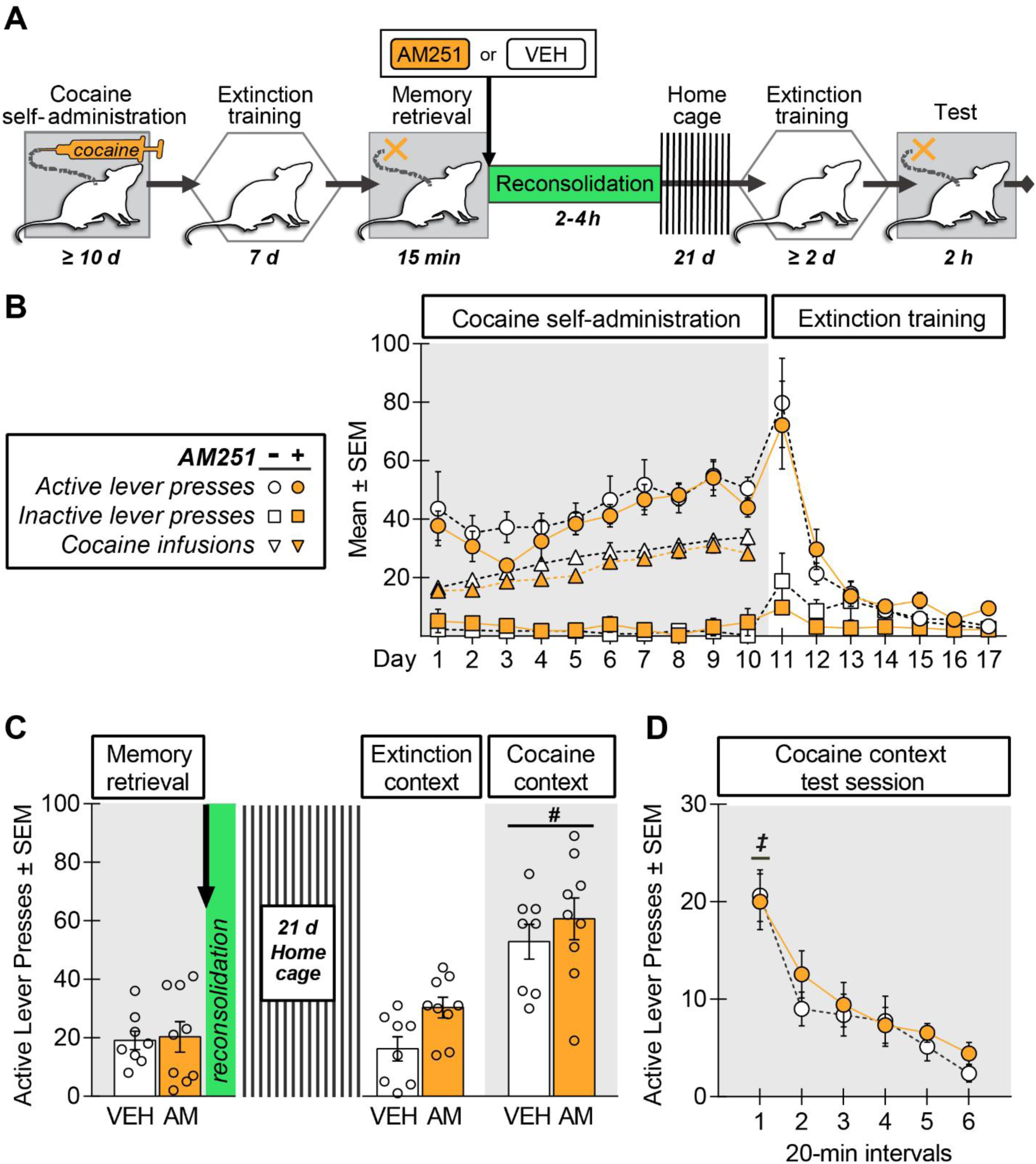
Systemic AM251 administration *during* memory reconsolidation does not alter drug context-induced cocaine-seeking behavior 24 days later. **(A)** Experimental timeline. The 15-min memory-retrieval session was immediately followed by systemic administration of AM251 (AM; 3 mg/kg, i.p.; *n* = 9) or VEH (*n* = 8). After 21 days of home cage stay, rats received at least two additional daily extinction sessions with ≤ 25 active-lever responses prior to the test of drug-seeking behavior in the cocaine-paired context. **(B)** Lever responses and cocaine infusions during drug self-administration and extinction training. **(C)** Active-lever responses during the memory-retrieval session in the cocaine-paired context before treatment (*arrow*) and after treatment followed by 21 days of home cage stay, upon first re-exposure to the extinction context and the cocaine-paired context at test. **(D)** Time course of active-lever responses in the cocaine-paired context at test. **Symbols**: ANOVA *^#^*context main effect, p < 0.05; *^‡^*time simple main effects (interval 1 > 2-6), Tukey’s tests, ps < 0.05.

### Experiment 4: Effects of memory reconsolidation and CB1R antagonism on BLA protein expression and phosphorylation

Experiment 4 evaluated memory reconsolidation-related changes in immediate-early gene expression and glutamate-receptor subunit expression and phosphorylation. The procedures in Experiment 4 were identical to those in Experiment 1 except that rats were euthanized 45 min after memory retrieval or no-memory retrieval (home-cage stay) and pharmacological treatment (**Fig. 4A**). This euthanasia time point was selected based on the activation kinetics of Arc and zif268 (Lee, 2004; Li et al., 2005) and associated changes in glutamate-receptor expression and phosphorylation in other models of synaptic plasticity (Clem and Huganir, 2010; Rao-Ruiz et al., 2011). The brains were flash frozen in isopentane and stored for analysis of BLA IEG and glutamate-receptor subunit expression and glutamate-receptor subunit phosphorylation using western blotting.

**Figure 4.**
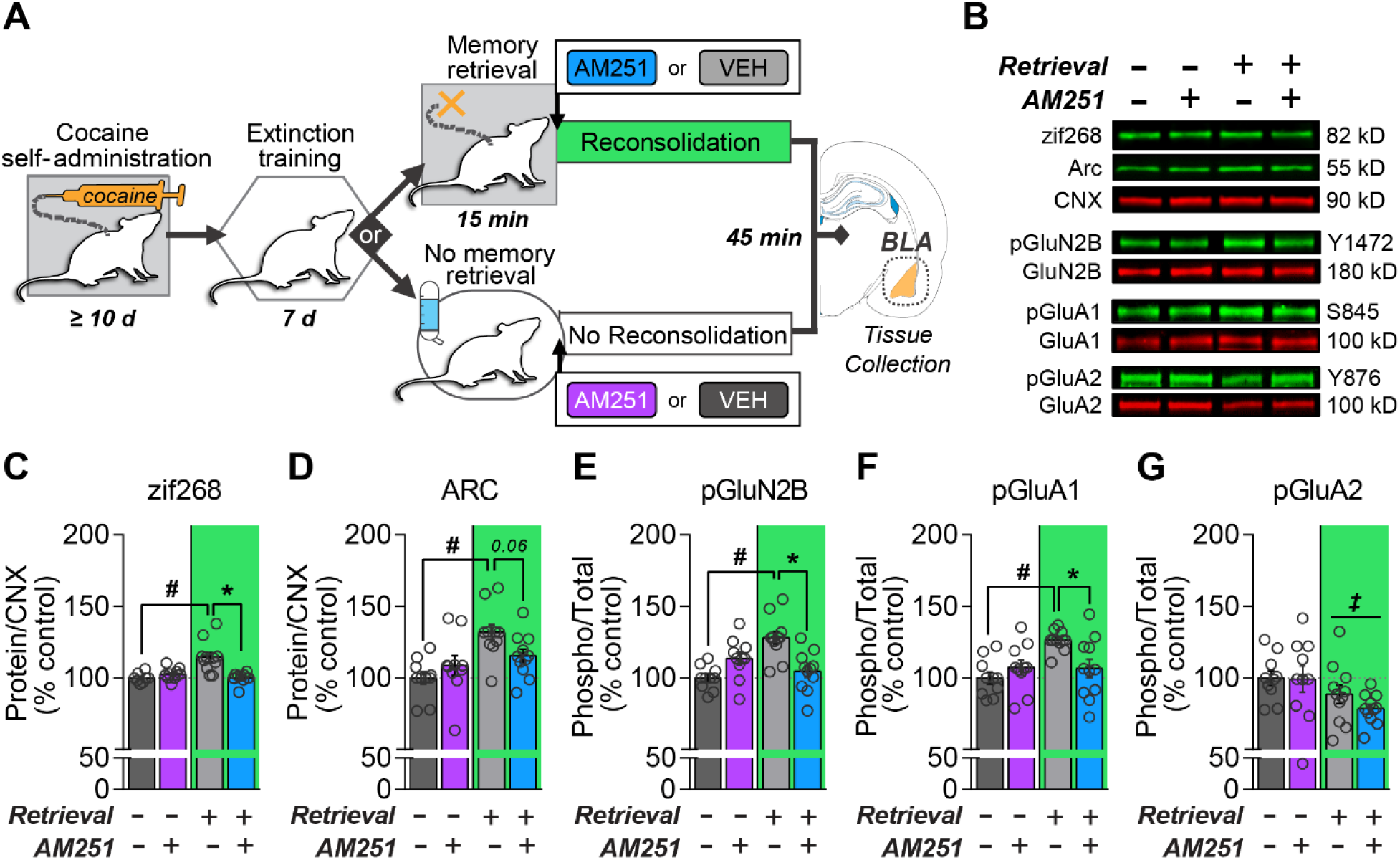
Systemic AM251 administration after memory retrieval prevents changes in BLA IEG expression and glutamate receptor subunit phosphorylation *during* memory reconsolidation. **(A)** Experimental timeline. Systemic AM251 (3 mg/kg, i.p.) or VEH was administered immediately after cocaine-memory retrieval or no-memory retrieval (*n* = 10 to 11 per group). BLA tissue samples were collected 45 min later for analysis of full tissue homogenates. **(B)** Representative western blots. Mean **(C)** zif268 and **(D)** ARC levels normalized to the loading control, calnexin (CNX). Mean levels of **(E)** GluN2B phosphorylation at Y1472 (pGluN2B), **(E)** GluA1 phosphorylation at S845 (pGluA1), and **(F)** GluA2 phosphorylation at Y876 (pGluA2) normalized to total protein levels. See extended **Figure 4-1** for control protein measures. Values are expressed as a percentage of the no-memory retrieval VEH control group. **Symbols**: ANOVA, *^#^*retrieval simple main effect, Sidak’s test, p < 0.05, **\***treatment simple main effect, Sidak’s test, p < 0.05, *^‡^*retrieval main effect, p < 0.05.

### Western Blotting

Brains were stored at −80 °C before the collection of BLA tissue punches (o.d. 0.75 mm). Punched tissue was stored at −80 °C in lysis buffer containing 10 mM HEPES, 1% SDS, and 1x protease and phosphatase inhibitor cocktails (Sigma Aldrich, St. Louis, MO). Samples were thawed, manually homogenized, and total tissue homogenate protein concentrations were determined using the Biorad detergent-compatible protein assay (R^2^ ≥ 0.99). After electrophoresis and transfer, membranes were dried overnight at 4°C. The next day, membranes were reactivated in methanol and blocked before incubation overnight in Odyssey blocking buffer (Li-Cor Biosciences, Lincoln, NE) with 0.2% Tween 20 and primary antibodies targeting Arc (Cat# sc-17839, RRID:AB_626696), zif268 (Cat# sc-189, RRID: AB_2231020), NMDAR subunit 2B (**GluN2B**; Cat# 06-600, RRID:AB_310193), phospho-Tyr^1472^-GluN2B (**pGluN2B**; Cat# p1516-1472, RRID:AB_2492182), GluA1 (Cat# sc-13152, RRID:AB_627932), phospho-Ser^845^-GluA1 (**pGluA1**; Cat# AB5849, RRID:AB_92079), GluA2 (Cat# MABN1189, RRID:AB_2737079), phospho-Tyr^876^-GluA2 (**pGluA2**; Cat# 4027, RRID:AB_1147622), calnexin (**CNX**; Cat# ADI-SPA-860, RRID:AB_10616095), or glyceraldehyde-3-phosphate dehydrogenase (**GAPDH**; Cat# ab8245, RRID:AB_2107448). Membranes were then washed and incubated for 1 h in Odyssey blocking buffer with 0.2% Tween 20 and 0.01% SDS with the following near-infrared fluorescent secondary antibodies: IRDye® 800CW goat anti-mouse (Cat# 926-32210, RRID:AB_621842), IRDye® 800CW goat anti-rabbit (Cat# 926-32211, RRID:AB_621843), IRDye® 680RD donkey anti-mouse (Cat# 926-68072, RRID:AB_10953628), and IRDye® 680LT goat anti-rabbit (Cat# 926-68021, RRID:AB_10706309). For multiplexed targets (based on antibody availability), the 800-nm channel was used to detect the lowest-abundance, phospho-proteins as it provides lower background, maximizing detection sensitivity (Schutz-Geschwender et al., 2004). Total proteins were detected in the 680-nm channel. Membranes were digitally imaged using a Li-Cor Odyssey CLX. Integrated optical intensity values for target protein bands were derived using Li-Cor Image Studio Software (RRID: SCR_015795). Total protein levels were normalized to a loading control and expressed as a percentage of the comparator group (Experiments 1-3: VEH-treated; Experiment 4: VEH-no-memory retrieval). Phospho-specific values were normalized to total protein values. A lane-normalization factor was calculated for the total and house-keeping proteins by dividing the lane’s integrated optical intensity value by the highest signal value on the blot. Target-protein integrated intensity values were divided by the respective loading control’s lane-normalization factor to normalize signal values across blots (Schutz-Geschwender et al., 2004).

### Experiment 5: Effects of post-retrieval CB1R antagonism on excitatory synaptic transmission in BLA PNs at memory reconsolidation

Experiment 5 examined the effects of memory reconsolidation and systemic CB1R antagonism on glutamatergic EPSCs in BLA PNs, the primary output neurons of the BLA (McDonald, 1984). The procedures in Experiment 5 were identical to those in Experiment 1 except that rats were exposed to the cocaine-paired context for 15 min (memory retrieval) or remained in their home cages (no-memory retrieval) before systemic AM251 (3 mg/kg, i.p.) or VEH administration (**Fig. 6A**). Rats were euthanized and brain slices were prepared for whole-cell electrophysiology recordings within an average of 30 min of treatment.

**Figure 5.**
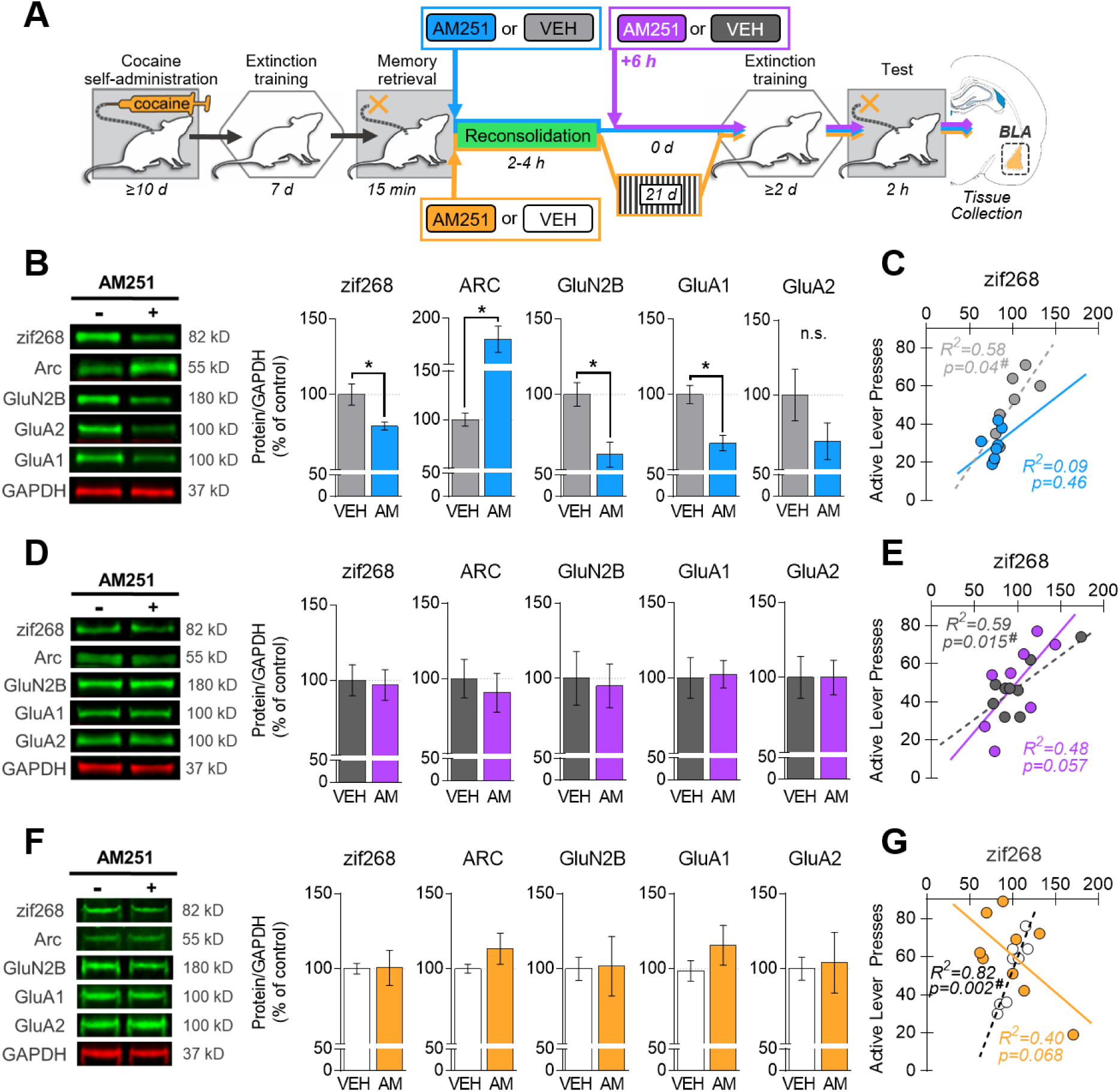
Systemic AM251 administration after memory retrieval elicits time-dependent changes in BLA IEG and total glutamate receptor subunit expression levels at test and in relationship with the magnitude of drug-seeking behavior. **(A)** Experimental timeline. Systemic AM251 (AM; 3 mg/kg, i.p.) or VEH was administered immediately or six hours after cocaine-memory retrieval. Cocaine-seeking behavior was assessed in the cocaine-paired context three or 24 days later, after at least two extinction sessions in Experiments 1-3. BLA tissue samples were collected immediately after the test session. **(B)** Effects of systemic AM251 administration immediately after memory retrieval on total protein levels (mean ± SEM) in the BLA at test, three days later in Experiment 1. **(C)** Relationship between total zif268 protein levels and active-lever presses at test in Experiment 1. **(D)** Effects of systemic AM251 administration 6 hours after memory retrieval on total protein levels (mean ± SEM) in the BLA at test, three days later in Experiment 2. **(E)** Relationship between total zif268 protein levels and active-lever presses at test in Experiment 2. **(F)** Effects of systemic AM251 administration immediately after memory retrieval on total protein levels (mean ± SEM) in the BLA at test, 24 days later in Experiment 3. **(G)** Relationship between total zif268 protein levels and active-lever presses at test in Experiment 3. Values were normalized to the loading control, GAPDH, and expressed as a percentage of the VEH-treated group. **Symbols**: *\**t-test, p < 0.05; **^#^**Pearson’s r correlation coefficient, p < 0.05 (*n* = 7-9 per group).

**Figure 6.**
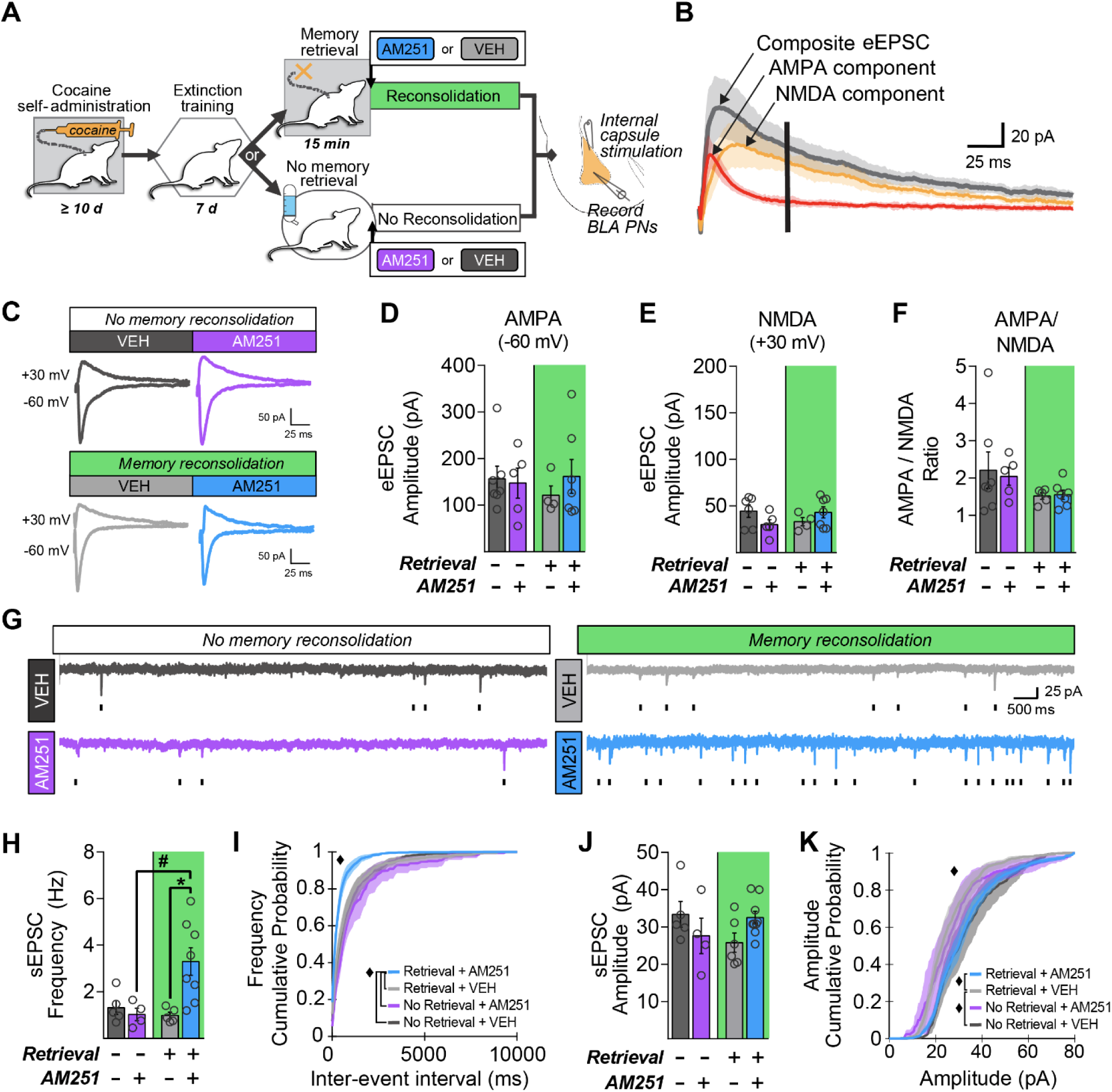
Systemic AM251 administration after memory retrieval increases sEPSC frequency and amplitude in BLA PNs during cocaine-memory reconsolidation. **(A)** Experimental timeline. AM251 (3 mg/kg, i.p.) or VEH was administered immediately after cocaine-memory retrieval or no-memory retrieval (See **Fig. 6-1** for behavioral history; *n* = 4 to 5 per group). Brain slices were prepared for whole-cell recording within an average of 30 min after treatment. Schematic depicting stimulating electrode placement in the internal capsule and recording electrode placement in the basal nucleus. **(B)** eEPSCs (mean across all cells recorded ± SEM) recorded at +30 mV in aCSF (*gray*) or AP5 (leaving the isolated AMPAR component, *red*), and superimposed digital subtraction (to visualize digitally-derived NMDAR component, *orange*). NMDAR-mediated eEPSC amplitudes were measured 50 ms after stimulus onset to eliminate AMPAR component contribution (*black bar*). **(C)** Representative recordings of eEPSC obtained from BLA PN cells in aCSF, in each group at −60 mV and +30 mV holding potentials. Peak amplitude of AMPA (mean ± SEM, at −60 mV; **D**) and NMDA (mean ± SEM, at +30 mV; **E**) components in aCSF. **(F)** AMPA/NMDA ratio (mean ± SEM). **(G)** Representative sEPSC recordings with ticks indicating individual events. **(H)** sEPSC frequency (mean ± SEM) and **(I)** the cumulative probability distribution of sEPSC inter-event intervals (mean ± SEM). **(J)** sEPSC amplitude (mean ± SEM) and **(K)** the cumulative probability distribution of sEPSC amplitudes (mean ± SEM). **Symbols**: ANOVA, *^#^*retrieval simple main effect, Sidak’s test, p < 0.05; *\**treatment simple main effect, Sidak’s test, p < 0.05; *^♦^*Kruskal-Wallis test, p < 0.05, Dunn’s test, p < 0.05.

### BLA Brain Slice Electrophysiology

All rats used for electrophysiological recordings were deeply anesthetized with isoflurane (3-5%). Rats were transcardially perfused with ice-cold artificial cerebrospinal fluid (**aCSF**), which contained 124 mM NaCl, 26 mM NaHCO_3_, 2.5 mM KCl, 2.5 mM CaCl_2_, 2 mM MgCl_2_, 1 mM NaH_2_PO_4_, 10 mM D-glucose, and 1 mM kynurenic acid, and was bubbled with 95% O_2_/5% CO_2_ (pH 7.4). After perfusion, the brain was rapidly removed and sliced coronally (225 µm) in ice-cold dissection buffer that contained 220 mM sucrose, 26 mM NaHCO_3_, 2 mM KCl, 0.5 mM CaCl_2_, 5 mM MgCl_2_, 1.25 mM NaH_2_PO_4_, 2 mM Na-pyruvate, 1 mM ascorbic acid, 10 mM D-glucose, 1 mM kynurenic acid, and was bubbled with 95% O_2_/5% CO_2_. Tissue slices containing the BLA were incubated in aCSF (with 1 mM kynurenic acid) at 35-37 ⁰C, after which they were incubated in room temperature aCSF (with 1 mM kynurenic acid) until used. Slices were discarded within 4 h of slice preparation.

Slices were transferred to a recording chamber and continually perfused (∼5 mL/min) in aCSF (without kynurenic acid) at a bath temperature of 32-35 °C. All pharmacological agents were dissolved in aCSF and were applied via bath perfusion. BLA pyramidal PNs were visually identified using differential interference contrast imaging through an Olympus 60x (0.9 NA) water-immersion objective. Whole-cell patch-clamp recordings were made using glass pipettes with a resistance of 2-3 MΩ when filled with internal solution that contained 130 mM CsCl, 4 mM NaCl, 0.5 mM CaCl_2_, 10 mM HEPES, 5 mM EGTA, 4 mM Mg-ATP, 0.5 mM Na_2_-GTP, 5 mM QX-314, 0.1 mM spermine, and 0.03 mM Alexa568 hydrazide dye with pH adjusted to 7.2–7.3 with CsOH. Cells were voltage-clamped at −60 or +30 mV. Signals were digitized at 20 kHz, low-pass filtered at 10 kHz, and additionally filtered at 2 kHz for presentation. To evoke EPSCs (**eEPSCs**), a concentric bipolar stimulating electrode was placed into the internal capsule (**IC**), and tissue was stimulated by current pulses (20-50 µA, 0.1-ms duration at a frequency of 0.1 Hz). Glutamatergic responses were pharmacologically isolated using 1 µM strychnine and 10 µM gabazine to block glycine and GABA_A_ receptors, respectively. AMPAR-mediated synaptic currents were isolated by adding 50 µM D-2-amino-5-phosphonovalerate (**AP5**), a broad-spectrum NMDAR antagonist. Ten eEPSCs were averaged in each condition, and their mean amplitudes were analyzed with pClamp software (RRID: SCR_011323). In the absence of AP5, NMDAR-mediated eEPSCs were measured as the average amplitude of the EPSC 50 ms after the onset of the stimulus (V_h_ = +30 mV), when AMPAR-mediated contributions are negligible (**Fig. 6B**). AMPA/NMDA ratios were calculated as the average inward peak current amplitude (V_h_ = −60 mV) divided by the outward current (V_h_ = +30 mV) amplitude at 50 ms after the onset of the stimulus. The rectification index was calculated as the average peak eEPSC amplitude at −60 mV divided by the average peak eEPSC amplitude at +30 mV, both recorded in the presence of AP5 to pharmacologically isolate the AMPAR component of the eEPSC. Spontaneous EPSCs (**sEPSCs**) from at least 300s long whole-cell recordings in each condition were analyzed with MiniAnalysis Program software. sEPSCs were detected automatically with an amplitude detection threshold of 2.5x the amplitude of the peak to peak of the noise. Events were visually confirmed as previously described (Richardson and Rossi, 2017), and the frequency and mean peak amplitude at −60 mV and +30 mV was measured.

### Experimental Design and Statistical Analysis

To identify potential pre-existing group differences in behavioral and drug history, active- and inactive-lever presses and cocaine intake during drug self-administration training (last 3 sessions) and non-reinforced lever presses during extinction training (first 7 extinction sessions) and during the memory-retrieval session were analyzed using separate mixed-factorial or univariate ANOVAs with subsequent treatment group as the between-subjects factor and time (session) as the within-subject factor, or between-subjects t-tests, where appropriate. Non-reinforced lever presses during the first post-treatment exposure to the extinction context and to the cocaine context were analyzed using mixed-factorial ANOVAs with memory retrieval (retrieval, no-memory retrieval) and treatment (AM251, VEH) as between-subjects factors and context (extinction, cocaine-paired) and time (20-min interval) as within-subjects factors, where appropriate. For Experiment 1-3, normalized total protein levels at test were analyzed using t-tests. For Experiments 4-5, phospho-protein levels, total protein levels, peak and mean eEPSC amplitudes, and sEPSC frequency at memory reconsolidation were analyzed using separate ANOVAs with treatment and memory retrieval as between-subjects factors. Significant interactions and main effects were further analyzed using Sidak’s or Tukey’s *post hoc* tests. Cumulative probability distributions of sEPSC amplitudes and inter-event intervals across groups (memory retrieval or no-memory retrieval, with AM251 or VEH) were analyzed using non-parametric Kruskal-Wallis tests with Dunn’s *post hoc* tests. The relationships between active-lever presses and total protein levels at test were analyzed using Pearson’s *r* correlational coefficients. Alpha was set at 0.05 for all analyses.

## RESULTS

### Behavioral and Drug History

There were no statistically significant differences between the groups in cocaine intake during drug self-administration training or in lever responding during drug self-administration training, extinction training, or memory retrieval in Experiments 1-5 (**Fig. 1-3; Extended Fig. 1-1, 2-1, 3-1, 6-1; Extended Tables 1-1, 2-1, 3-1, 4-1, 6-1)**.

### Experiment 1: Systemic CB1R antagonism during memory reconsolidation attenuates subsequent drug context-induced cocaine seeking

Systemic AM251 administration immediately after the 15-min cocaine-memory retrieval session (i.e., at the onset of memory reconsolidation) attenuated cocaine-seeking behavior at test in a context-dependent manner (**Fig. 1C**; 2 x 2 ANOVA context x treatment interaction, F_(1,13)_ = 10.93, p = 0.006; context main effect, F_(1,13)_ = 111.80, p = 0.0001; treatment main effect, F_(1,13)_ = 8.95, p = 0.01). Active-lever responding in the cocaine-paired context was greater than in the extinction context (Sidak’s test, p = 0.05). Furthermore, AM251 administered immediately after memory retrieval attenuated active-lever responding in the cocaine-paired (Sidak’s test, p = 0.05), but not the extinction, context relative to VEH. Time-course analysis of active-lever presses in the cocaine-paired context revealed that responding declined over time, and AM251 reduced responding during the first 20-min interval relative to VEH (**Fig. 1D**; 2 x 6 ANOVA treatment x time interaction, F_(5,65)_ = 4.11, p = 0.003, Tukey’s tests, p < 0.05; time main effect, F_(5,65)_ = 14.59, p = 0.0001; treatment main effect F_(1,13)_ = 11.86, p = 0.004). Inactive-lever responding remained low in both contexts independent of treatment (**Fig. 1-1**).

### Experiment 2: Systemic CB1R antagonism outside of the memory reconsolidation window does not alter subsequent drug context-induced cocaine seeking

Systemic AM251 administration 6 h after cocaine-memory retrieval (i.e., after reconsolidation into long-term memory stores) did not alter subsequent cocaine-seeking behavior relative to VEH (**Fig. 2C**). Active-lever responding in the cocaine-paired context was higher than in the extinction context regardless of treatment (2 x 2 ANOVA context main effect, F_(1,15)_ = 89.43, p < 0.0001), and delayed AM251 administration did not alter responding in either context relative to VEH (ANOVA treatment main and interaction effects, F_(1,15)_ ≤ 0.21, p ≥ 0.65). The time-course analysis of active-lever presses in the cocaine-paired context indicated that responding declined after the first 20-min interval independent of treatment (**Fig. 2D**; 2 x 6 ANOVA time main effect, F_(5,75)_ = 20.45, p < 0.0001, interval 1 > intervals 2-6, Tukey’s tests, p < 0.05; treatment main and interaction effects, Fs _(5,75)_ ≤ 0.88, p ≥ 0.50). Inactive-lever responding remained low in both contexts independent of treatment (**Fig. 2-1**).

### Experiment 3: Systemic CB1R antagonism during memory reconsolidation fails to alter drug context-induced cocaine seeking 24 days later

Systemic AM251 administration immediately after cocaine-memory retrieval (i.e., at the onset of memory reconsolidation) failed to alter cocaine-seeking behavior after a 21-d drug-free period followed by at least two extinction sessions later, relative to VEH (**Fig. 3C**). Active-lever responding in the cocaine-paired context was higher than in the extinction context regardless of treatment (2 x 2 ANOVA context main effect, F_(1,15)_ = 48.56, p < 0.0001; treatment main and interaction effects, Fs_(1,15)_ ≤ 3.32, p ≥ 0.09). Time-course analysis of active-lever presses in the cocaine-paired context indicated that responding declined after the first 20-min interval independent of treatment (**Fig. 3D**; 2 x 6 ANOVA time main effect, F _(5,75)_ = 20.08, p < 0.0001, interval 1 > intervals 2-6, Tukey’s tests, p < 0.05; treatment main and interaction effects, Fs _(5,75)_ ≤ 0.57, p ≥ 0.46). Inactive-lever responding remained low in both contexts independent of treatment (**Fig. 3-1**).

### Experiment 4: Systemic CB1R antagonism inhibits memory retrieval-induced molecular changes in the BLA during memory reconsolidation

In brain tissue collected during memory reconsolidation (**Fig. 4A-B**), BLA IEG expression varied as a function of memory retrieval and AM251 treatment. Memory retrieval increased zif268 expression relative to no-memory retrieval (i.e., home cage stay; **Fig. 4C**; 2 x 2 ANOVA, retrieval x treatment interaction, F_(1,38)_ = 18.91, p < 0.0001; retrieval main effect, F_(1,38)_ = 9.16, p = 0.004; treatment main effect, F_(1,38)_ = 9.08, p = 0.005). Systemic AM251 administration after memory retrieval reduced zif268 expression relative to VEH (Sidak’s test, p < 0.05), such that zif268 expression no longer differed from those in the no-memory retrieval controls. Similar to zif268, memory retrieval increased Arc expression relative to no-memory retrieval (**Fig. 4D**; 2 x 2 ANOVA, retrieval x treatment interaction, F_(1,38)_ = 5.61, p = 0.02; retrieval main effect, F_(1,38)_ = 13.03, p = 0.0009; treatment main effect, F_(1,38)_ = 0.53, p = 0.47). Furthermore, systemic AM251 administration after memory retrieval modestly attenuated Arc expression during memory reconsolidation relative to VEH (Sidak’s test, p = 0.06), such that Arc expression no longer differed from those in the no-memory retrieval controls.

Similar to IEG expression, glutamate receptor subunit phosphorylation varied as a function of memory retrieval and AM251 treatment. These results are reported in **Fig 4E-G**.

Src-mediated phosphorylation of NMDAR GluN2B^Y1472^ (**pGluN2B**) facilitates proper GluN2B synaptic localization, learning, and amygdalar synaptic plasticity (Nakazawa et al., 2006). BLA pGluN2B levels varied as a function of memory retrieval and treatment (**Fig. 4F**). Memory retrieval increased pGluN2B relative to no-memory retrieval (retrieval x treatment interaction, F_(1,38)_ = 20.74, p < 0.0001; retrieval main effect, F_(1,38)_ = 5.60, p = 0.02; treatment main effect, F_(1,38)_ = 1.43, p = 0.24). Systemic AM251 administration after memory retrieval reduced pGluN2B during memory reconsolidation relative to VEH (Sidak’s test, p < 0.05), such that pGluN2B levels no longer differed from those in the no-memory retrieval controls. Notably, a trend for an AM251-induced increase in total GluN2B levels (**Fig. 4-1B**; treatment main effect, F_(1,38)_ = 8.40, p = 0.006; retrieval main and interaction effects, Fs_(1,38)_ ≤ 3.72, p ≥ 0.06) could have enhanced this effect by increasing the denominator.

PKA-mediated phosphorylation of AMPAR GluA1^S845^ (**pGluA1**) promotes GluA1 trafficking to the postsynaptic density and fear-memory destabilization after memory retrieval (Clem and Huganir, 2010). BLA pGluA1 levels varied as a function of memory retrieval and AM251 treatment (**Fig. 4F**; retrieval x treatment interaction, F_(1,38)_ = 8.04, p = 0.007; retrieval main effect, F_(1,38)_ = 7.15, p = 0.01; treatment main effect, F_(1,38)_ = 1.70, p = 0.20) with no change in total GluA1 levels (**Fig. 4-1C**; all retrieval and treatment main and interaction effects, Fs_(1,38)_ ≤ 1.83, p ≥ 0.18). Memory retrieval increased pGluA1 relative to no-memory retrieval (Sidak’s test, p < 0.05). Moreover, systemic AM251 administration after memory retrieval reduced pGluA1 during memory reconsolidation relative to VEH (Sidak’s test, p < 0.05), such that pGluA1 levels no longer differed from those in no-memory retrieval controls.

Src-mediated phosphorylation of AMPAR GluA2^Y876^ (**pGluA2**) disrupts GluA2 association with postsynaptic density scaffolding proteins, thereby reducing GluA2 synaptic expression (Hayashi and Huganir, 2004). Memory retrieval reduced BLA pGluA2 during memory reconsolidation relative to no-memory retrieval (**Fig. 4G**; retrieval main effect, F_(1,38)_ = 6.78, p = 0.01; all treatment main and interaction effects, Fs_(1,38)_ ≤ 0.78, p ≥ 0.38), without altering total GluA2 levels (**Fig. 4-1D**; all Fs_(1,38)_ < 1.59, p > 0.22). In contrast, AM251 failed to alter pGluA2 or total GluA2 levels.

### Experiments 1-3: Systemic CB1R antagonism during memory reconsolidation inhibits molecular changes in the BLA during reinstatement three days, but not 24 days, post treatment

To capture potential protracted effects of AM251 on protein expression during the reinstatement test, BLA tissue was collected immediately after the 2-h test session in the cocaine-paired context in Experiments 1-3 (**Fig. 5A**). In Experiment 1, systemic AM251 administration immediately after memory retrieval (i.e., during memory reconsolidation) significantly reduced BLA zif268 (t _(13)_ = 2.94, p = 0.01), total GluN2B (t _(13)_ = 3.48, p = 0.004), and total GluA1 (t _(13)_ = 4.41, p = 0.001) expression relative to VEH three days post treatment. AM251 administration also increased Arc expression (t _(13)_ = 4.61, p = 0.0005) and failed to alter total GluA2 levels (t _(13)_ = 1.63, p = 0.125) relative to VEH at the same time point (**Fig. 5B**). Furthermore, there was a positive correlation between zif268 expression and active-lever responding at test three days after VEH treatment (**Fig. 5C**; *r* = 0.82, p = 0.05). This correlation was not observed after AM251 treatment (*r* = 0.31, p = 0.46) or between active-lever responding and other protein targets (**Table 5-1,** *r* ≤ 0.40, p ≥ 0.12).

In Experiment 2, systemic AM251 administration 6 h after memory retrieval (i.e., outside the memory reconsolidation window) failed to alter BLA zif268, Arc, GluN2B, GluA1, or GluA2 levels relative to VEH at test, three days post treatment (**Fig. 5D**; t_(13)_ ≤ 0.51, p ≥ 0.61). Furthermore, there was a positive correlation between BLA zif268 expression and active-lever responding at test three days after VEH treatment (*r* = 0.77, p = 0.02) and a trend for a similar positive correlation after AM251 treatment (**Fig. 5E**; *r* = 0.69, p = 0.058).

In Experiment 3, systemic AM251 administration immediately after memory retrieval did not alter zif268, Arc, GluN2B, GluA1, or GluA2 levels relative to VEH at test, 24 days later (**Fig. 5F**; all t_(15)_ ≤ 1.18, p ≥ 0.26). However, there was a positive correlation between BLA zif268 expression and active-lever responding at test after memory retrieval and VEH treatment (**Fig. 5G**; *r* = 0.90, p = 0.002) and a trend for a negative correlation following AM251 treatment (*r* = - 0.63, p = 0.07).

### Experiment 5: Systemic CB1R antagonism after memory retrieval increases sEPSC frequency in BLA PNs

To determine if AM251-induced memory reconsolidation impairments and associated molecular changes were paralleled by changes in synaptic transmission in the BLA, whole cell patch-clamp recordings were obtained from BLA PNs in slices prepared on average 30 min after memory retrieval or no-memory retrieval and systemic AM251 or VEH treatment (**Fig. 6A**). Application of the NMDAR antagonist, APV (50 µM), was used to pharmacologically isolate the AMPAR component, and subsequent digital subtraction revealed the NMDA component, of the composite eEPSC (**Fig. 6B**). The AMPAR and NMDAR components were operationalized as the peak amplitude and the amplitude at 50 ms of the composite response, respectively. Neither memory retrieval nor AM251 altered the AMPAR-mediated eEPSC amplitude (**Fig. 6C-D**; ANOVA, all retrieval and treatment Fs_(1,19)_ ≤ 0.57, p ≥ 0.46), NMDAR-mediated eEPSC amplitude (**Fig. 6E**; ANOVA, all Fs_(1,18)_ ≤ 3.89, p ≥ 0.06), or the AMPA/NMDA eEPSC amplitude ratio (**Fig. 6F**; ANOVA, all Fs_(1,17)_ ≤ 2.73, p ≥ 0.12).

In contrast to the lack of effects on eEPSCs, the sEPSC frequency varied as a function of memory retrieval and systemic AM251 treatment (**Fig. 6G**). Memory retrieval followed by VEH did not alter the mean sEPSC frequency relative to no-memory retrieval (ANOVA, retrieval x treatment interaction, F_(1,18)_ = 6.95, p = 0.02; all other Fs_(1,18)_ ≤ 4.17, p ≥ 0.06). However, AM251 administration after memory retrieval, but not after no-memory retrieval, increased the mean sEPSC frequency relative to VEH (**Fig. 6H**; Sidak’s test, p < 0.05) and produced a leftward shift in the cumulative probability distribution of sEPSC inter-event intervals relative to VEH after memory retrieval or AM251 after no-memory retrieval (**Fig. 6I**; Kruskal-Wallis test, H_(3)_ = 72.14, p < 0.0001, Dunn’s test, p < 0.05). The effect of memory retrieval on sEPSC amplitude was also dependent on memory retrieval and treatment (**Fig. 6J**; ANOVA, context x treatment interaction, F _(1,20)_ = 4.737, p = 0.04; all other Fs _(1,20)_ ≤ 0.22, p ≥ 0.65), but *post-hoc* comparisons did not indicate group differences. Memory retrieval followed by VEH produced a leftward shift in the cumulative probability distribution of sEPSC amplitudes relative to no-memory retrieval (Kruskal-Wallis test, H_(3)_ = 11.67, p = 0.01, Dunn’s test, p < 0.05), and AM251 administration after memory retrieval inhibited this shift relative to VEH (**Fig. 6K**; Dunn’s test, p < 0.05).

To verify our measurements of non-pharmacologically separated composite EPSCs, we repeated our measurements on pharmacologically isolated AMPAR eEPSCs (**Fig. 7A**). In the presence of the NMDAR antagonist, AP5 (50 µM), neither memory retrieval nor systemic AM251 administration altered synaptic responses to IC stimulation, including peak AMPAR eEPSC amplitude at −60 mV (**Fig. 7B**; ANOVA, all Fs_(1,35)_ ≤ 3.30, p ≥ 0.08) and +30 mV (**Fig. 7C**; ANOVA, all Fs_(1,35)_ ≤ 0.68, p ≥ 0.42), or the AMPAR rectification index (**Fig. 7D**; ANOVA, all Fs_(1,34)_ ≤ 0.11, p ≥ 0.75).

**Figure 7.**
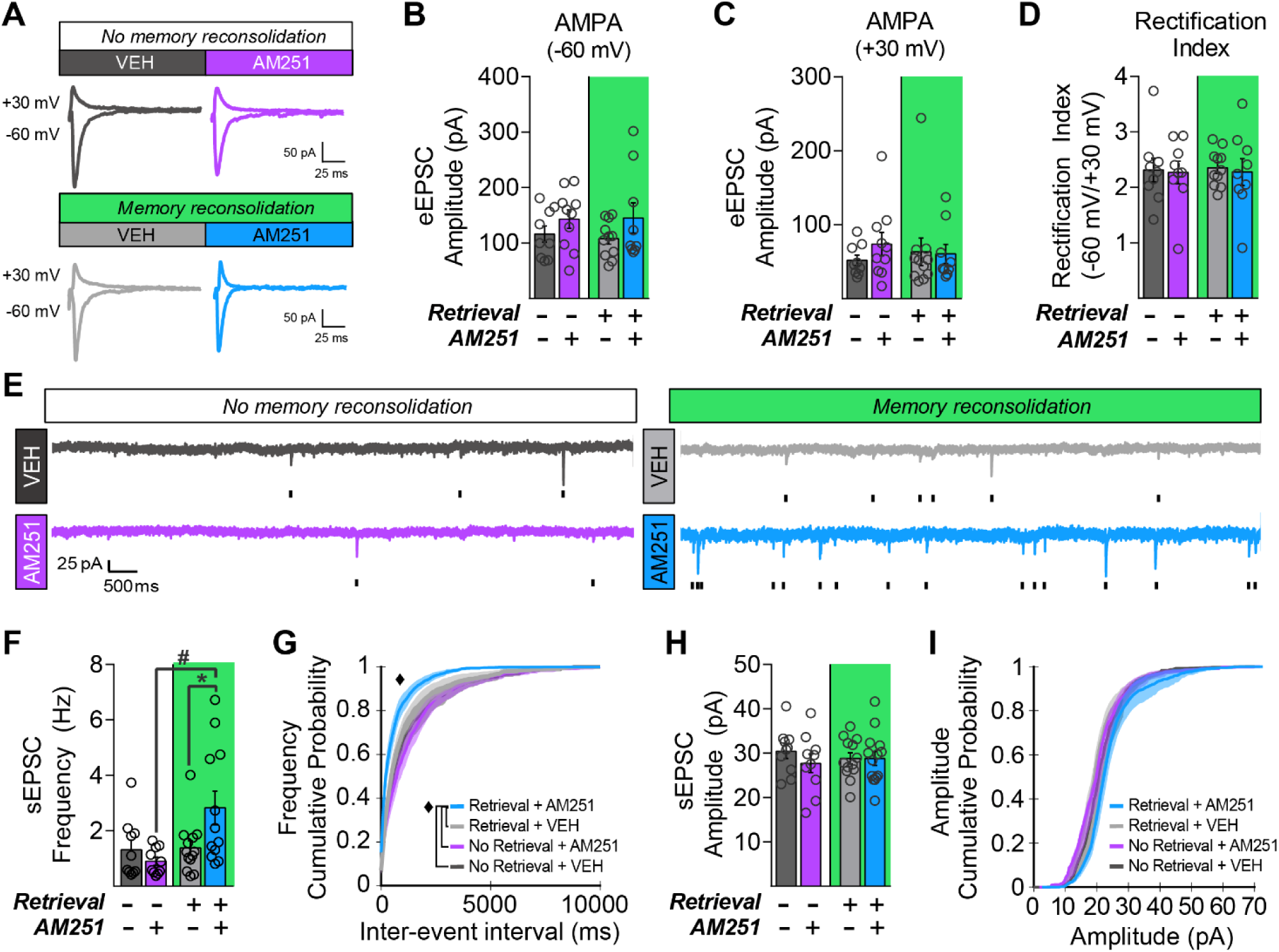
Systemic AM251 administration after memory retrieval increases AMPAR-mediated sEPSC frequency in BLA PNs during cocaine-memory reconsolidation. **(A)** Representative eEPSCs recordings in the presence of the NMDAR antagonist, AP5, at +30 mV and −60 mV. Peak AMPAR-mediated eEPSC amplitudes (mean ± SEM) **(B)** at – 60 mV and **(C)** +30 mV. **(D)** Rectification index calculated as the ratio of the peak eEPSC amplitude at −60 mV and +30 mV in the presence of AP5. **(E)** Representative sEPSC recordings with ticks indicating individual events. **(F)** sEPSC frequency (mean ± SEM). **(G)** Cumulative probability distribution of sEPSC inter-event intervals (mean ± SEM). **(H)** sEPSC amplitude (mean ± SEM) and **(I)** cumulative probability distribution of amplitudes (mean ± SEM). **Symbols**: ANOVA, *^#^*retrieval simple main effect, Sidak’s test, p < 0.05; *\**ANOVA, treatment simple main effect, Sidak’s test, p < 0.05; *^♦^*Kruskal-Wallis test, p < 0.05, Dunn’s test, p < 0.05.

In the presence of AP5, the mean AMPAR sEPSC frequency varied as a function of memory retrieval and AM251 treatment (**Fig. 7E**). Memory retrieval followed by VEH did not alter mean sEPSC frequency relative to no-memory retrieval (ANOVA, retrieval x treatment interaction, F_(1,41)_ = 5.53, p = 0.02; retrieval main effect, F_(1,41)_ = 6.24, p = 0.02; treatment main effect, F_(1,41)_ = 1.68, p = 0.20). However, systemic AM251 administration after memory retrieval, but not after no-memory retrieval, increased sEPSC frequency relative to VEH (**Fig. 7F**; Sidak’s test, p < 0.05) and produced a leftward shift in the cumulative probability distribution of sEPSC inter-event intervals relative to VEH after memory retrieval or AM251 after no-memory retrieval (**Fig. 7G**; Kruskal-Wallis test, H_(3)_ = 43.18, p < 0.0001; Dunn’s test, p < 0.05). Neither memory retrieval nor AM251 treatment altered the mean amplitude of sEPSCs (**Fig. 7H**; ANOVA, all Fs_(1,43)_ ≤ 0.73, p ≥ 0.41, Fig. figure caption) or the cumulative probability distribution of sEPSC amplitudes (**Fig. 7I**; Kruskal-Wallis test, H _(3)_ = 0.28, p = 0.96).

## DISCUSSION

The main finding of this study is that CB1R signaling critically modulates memory reconsolidation processes necessary for subsequent drug context-induced cocaine-seeking behavior in an instrumental model of drug relapse. Furthermore, memory retrieval induces CB1R-dependent changes in IEG expression, glutamatergic receptor subunit phosphorylation, and excitatory synaptic transmission in the BLA during memory reconsolidation, and some of these changes are predictive of the magnitude of subsequent drug-seeking behavior.

Systemic CB1R antagonism during cocaine-memory reconsolidation (i.e., immediately after memory retrieval) reduced drug context-induced cocaine-seeking behavior three days later, relative to VEH (**Fig. 1**). The CB1R antagonist, AM251 does not alter inhibitory avoidance (Gobira et al., 2013) or grooming behaviors (Hodge et al., 2008) at similar doses, suggesting that AM251 is not acutely aversive. Furthermore, CB1R antagonism alone did not alter the expression of drug-seeking behavior despite its long half-life (i.e., 22-h; McLaughlin et al., 2003) (**Fig. 2**). These observations suggest that CB1R signaling is necessary for cocaine-memory reconsolidation in an instrumental model of drug relapse, thereby expanding on the known involvement of CB1Rs in Pavlovian morphine, nicotine, and methamphetamine memory reconsolidation (Yu et al., 2009; Fang et al., 2011; De Carvalho et al., 2014). CB1R signaling is not required for social reward memory reconsolidation (Achterberg et al., 2014); as such, selective effects of CB1R antagonism on memory reconsolidation across drug classes and paradigms are especially encouraging from a substance use disorder treatment perspective.

Systemic CB1R antagonism inhibited molecular adaptations in the BLA during memory reconsolidation and at the time of the test for drug context-induced cocaine-seeking behavior three days later. Specifically, memory retrieval augmented zif268 and Arc expression during memory reconsolidation, and this effect was blocked by AM251 (**Fig. 4**). CB1R-dependent IEG expression in the BLA is probably required for memory re-stabilization given that intra-BLA zif268 or Arc antisense administration disrupts cocaine-CPP and explicit conditioned stimulus (CS)- cocaine-memory reconsolidation in other paradigms (Lee et al., 2005, 2006; Theberge et al., 2010; Alaghband et al., 2014). At test, there was a CB1R-dependent, positive relationship between the extent of BLA zif268 expression and the magnitude of drug context-induced cocaine-seeking behavior (**Fig. 5C**). This finding is consistent with reports of increased zif268 mRNA expression during reinstatement (Hearing et al., 2010; Ziółkowska et al., 2011) and suggests that BLA zif268 expression tracks drug context-induced motivation for cocaine. However, the *memory retrieval-dependent* AM251 effects indicate that diminished motivation for cocaine at test is due to CB1R antagonist-induced impairment in cocaine-memory strength or retrieval link establishment during memory reconsolidation.

Interestingly, cocaine-memory retrieval triggered CB1R-dependent increases in pGluN2B^Y1472^ and pGluA1^S845^, consistent with enhanced GluN2B synaptic stability (Nakazawa et al., 2006) and GluA1 synaptic recruitment (Hong et al. 2010), respectively, as well as CB1R-independent decreases in pGluA2^Y876^ (**Fig. 4**). Previous research has indicated the necessity of PKA activation (Arguello et al. 2014) and the importance of NMDAR-dependent CI-AMPAR (GluA2-containing) endocytosis followed closely by CP-AMPAR (GluA2-lacking) synaptic trafficking in the amygdala for memory reconsolidation (Clem and Huganir, 2010; Hong et al., 2013; Lopez et al., 2015; Yu et al., 2016). In our study, decreases in pGluA2^Y876^, consistent with diminished Src-mediated CI-AMPAR endocytosis (Hayashi and Huganir, 2004), likely captured a time point when CI-AMPAR synaptic expression was restored after transient endocytosis, while CP-AMPAR synaptic insertion was still ongoing (Rao-Ruiz et al. 2015). Thus, cocaine-memory reconsolidation may involve CB1R- and NMDAR-dependent CP-AMPAR synaptic insertion and CB1R-independent CI-AMPAR endocytosis.

The robust effects of memory retrieval on IEG expression and glutamate-receptor subunit phosphorylation were paralleled by subtle influences on BLA synaptic physiology observed in slices prepared on average 30 min after memory retrieval and systemic AM251 treatment. During reconsolidation, IC stimulation did not reveal changes in AMPA/NMDA ratio, eEPSC peak amplitude, or eEPSC rectification in BLA PNs (**Fig. 6-7**). Similarly, fear-memory reconsolidation is associated with equal-conductance exchange of CI-AMPARs to CP-AMPARs and no change in IC→BLA synaptic strength, with a transient increase in rectification reported in slices prepared 5 min, but not one hour, after memory retrieval (Hong et al. 2013; Rao-Ruiz et al. 2015). Since we observed molecular changes in the BLA 45 min after memory retrieval (**Fig. 4**), it is possible that the temporal dynamics of IC→BLA synaptic plasticity during cocaine-memory and fear-memory reconsolidation differ, or the timing of our synaptic physiology protocols did not permit the detection of transient changes in rectification. Finally, the observed molecular adaptations in the BLA might reflect changes in receptor trafficking at *non*-IC→BLA synapses or other cell types (i.e., GABAergic interneurons).

Importantly, we discovered memory retrieval-dependent increases in sEPSC frequency and reductions in peak sEPSC amplitude in BLA PNs (selected based on their responsiveness to IC stimulation, **Fig. 6-7**). Moreover, sEPSC frequency increases were only observed when CB1Rs were blocked. This suggests that potential cocaine-memory retrieval-associated plasticity occurs at *non*-IC excitatory inputs to BLA PNs (e.g., sensory cortices, prefrontal cortex, or ventral hippocampus; LeDoux, 2007; Sah et al., 2003), and systemic CB1R antagonism enables potentiation at these inputs, and may create “synaptic noise” that impedes plasticity associated with memory strength or reconsolidation. Thus, CB1R signaling may facilitate memory reconsolidation by reducing “synaptic noise.” Additionally, memory retrieval-dependent reductions in sEPSC peak amplitude (**Fig. 6J-K)** were observed in a CB1R- and NMDAR-dependent manner **(Fig. 7H-I**). If the rising phase of the NMDAR component contributed to the peak composite sEPSC, then increases in BLA pGluN2B^Y1472^ (**Fig. 4**) may explain the reductions in sEPSC amplitude given the lower open probability of GluN2B-containing NMDARs (Paoletti et al., 2013). The involvement of GluN2B in appetitive memory reconsolidation has not been investigated until now, but earlier studies indicate that GluN2B mediates Pavlovian fear-memory destabilization, but not re-stabilization (Ben Mamou et al., 2006; Milton et al., 2013). Consequently, future research will need to evaluate whether the contributions of GluN2B-containing NMDARs to memory reconsolidation are paradigm specific.

Collectively, cocaine-memory reconsolidation involves at least two forms of plasticity at BLA PN synapses: (a) a memory retrieval-induced increase in excitability or vesicle release probability of glutamatergic afferents that is inhibited by CB1R signaling and (b) an NMDAR- and CB1R-dependent decrease in sEPSC amplitude. The former finding implies that CB1R signaling is necessary to *reduce* glutamatergic synaptic excitation of BLA PNs during memory reconsolidation. Initially, this appears to be contradictory to effects of CB1R antagonism on memory reconsolidation in the present study and to the known role of glutamate in memory reconsolidation (Milton et al., 2008; Rao-Ruiz et al., 2015). However, it is consistent with the recognized role of CP-AMPARs in synaptic scaling, a plasticity mechanism important for rebalancing neuronal excitability (Diering and Huganir, 2018). CP-AMPARs preferentially accumulate at synapses after LTP (Turrigiano, 2008; Turrigiano et al., 2014; Diering and Huganir, 2018) and facilitate synaptic resistance to down-scaling and ubiquitination (Diering et al., 2014; Sanderson et al., 2016). Future studies will need to investigate whether synaptic scaling is a plasticity mechanism of memory reconsolidation.

Contrary to our hypothesis, the effects of AM251 on drug-seeking behavior and BLA protein expression were not present 24 days post treatment (**Fig. 5**). Transient amnesia could result from AM251-induced enhancement in extinction memory consolidation followed by spontaneous recovery. However, this was unlikely because brief re-exposure to the cocaine-paired context did not elicit either appreciable behavioral extinction or synaptic depression in IC→BLA synapses, a phenomenon associated memory extinction (Clem and Huganir, 2010; Rich et al., 2019). Furthermore, testing occurred immediately after two extinction sessions, as opposed to after time away from the testing environment that is required for spontaneous recovery (Rescorla, 2004). It is more likely, instead, that the transient effects of AM251 reflected the delayed availability of alternate memory traces following amnesia due to reconsolidation interference (Nadel and Moscovitch, 1997). Alternatively, AM251 could inhibit neural ensembles that form “retrieval links” (Lewis et al., 1968). Consistent with this idea, reactivation of neural ensembles that are active during encoding is necessary and sufficient for memory expression (Ramirez et al., 2013; Roy et al., 2016; Richards and Frankland, 2017).

In conclusion, systemic administration of AM251 in conjunction with re-exposure to the cocaine-paired context, a manipulation analogous to exposure therapy, blocks glutamatergic mechanisms associated with memory reconsolidation and transiently alleviates drug context-induced motivation to seek cocaine in a rodent model of drug relapse. Thus, AM251 may provide short-term benefit for individuals with cocaine-use disorders, similar to β-adrenergic receptor antagonism (Saladin et al., 2013). Furthermore, the CB1R may be a suitable therapeutic target for interference with the reconsolidation of maladaptive memories, even though AM251 also has some efficacy as a GPR55 (CB1 orphan receptor) agonist (Kapur et al., 2009) and µ-opioid receptors antagonist (Seely et al., 2012). While additional research is needed to determine the exact mechanisms of action for AM251, the present findings are interesting from a pharmacotherapeutic perspective. *Chronic* daily treatment with the CB1R inverse agonist, rimonabant, is effective for smoking cessation but produces detrimental side effects (Moreira and Crippa, 2009). Therefore, future studies will need to examine whether a less extensive regimen of *repeated* post-retrieval administration of AM251 or other CB1R antagonists can safely produce lasting reductions in cue reactivity and cocaine seeking without detrimental side effects.

## Acknowledgements

This research was funded by NIDA R01 DA025646 (RAF), NIDA F31 DA 045430 (JAH), NIAAA R01 AA026078 (DJR), NINDS T32 NS007431 (MAP), and Washington State Initiative 171 (JAH) and 502 (RAF) funds administered through the Washington State University Alcohol and Drug Abuse Research Program.

## EXTENDED DATA

**Figure 1-1.**
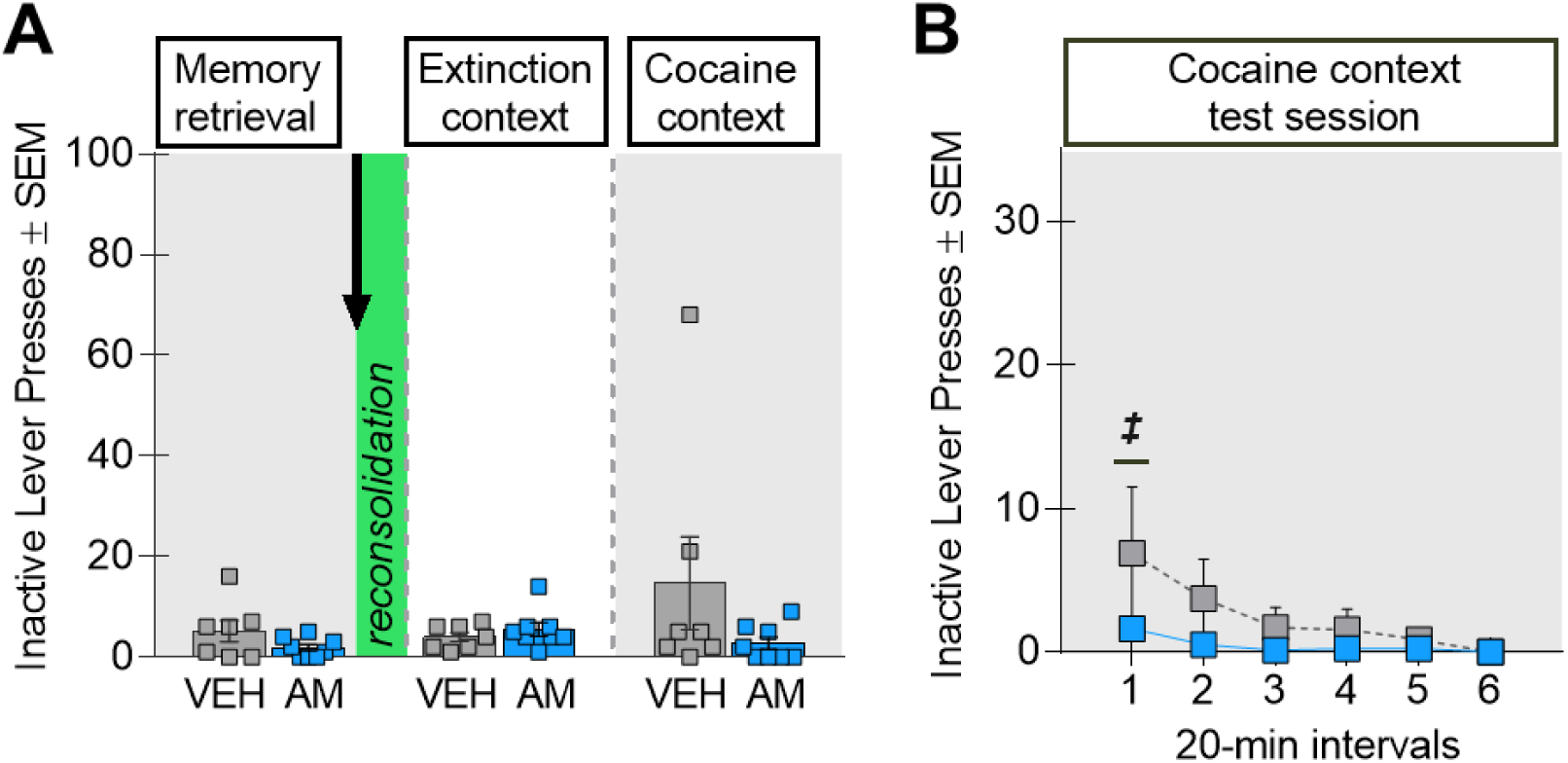
Inactive-lever responding in Experiment 1. **(A)** Inactive-lever responding was not different between the groups during the 15-min memory-retrieval session (*left panel*) before administration of AM251 (AM, *n* = 8) or VEH (*n* = 7) (t _(13)_ = 1.54, p = 0.15) (*arrow*). After treatment, inactive-lever responding remained low in the extinction and cocaine-paired contexts independent of treatment (ANOVA, all Fs_(1,13)_ ≤ 2.52, p ≥ 0.14). **(B)** Inactive-lever responding in the cocaine-paired context at test declined over time independent of treatment (ANOVA, time main effect, F_(5,65)_ = 2.71, p = 0.03, ***^‡^***time simple main effects, Tukey’s tests, interval 1 > 2-6, ps < 0.05).

**Table 1-1.** Experiment 1 Behavioral History

| Phase | Measure | Treatment Main Effects |  |  |  | Day Main Effects |  |  |  | Treatment x Day |  |  |  |
| --- | --- | --- | --- | --- | --- | --- | --- | --- | --- | --- | --- | --- | --- |
|  |  | Test | df | Statistic | p | Test | df | Statistic | p | Test | df | Statistic | p |
| Self-Administration | Active Lever | F | 1,13 | 0.60 | 0.45 | F | 9,117 | 0.86 | 0.57 | F | 9,117 | 0.29 | 0.74 |
|  | Inactive Lever | F | 1,13 | 2.29 | 0.15 | F | 9,117 | 1.16 | 0.33 | F | 9,117 | 1.20 | 0.30 |
|  | Cocaine Infusion | F | 1,13 | 0.15 | 0.71 | F | 9,117 | 20.19 | <0.0001 | F | 9,117 | 2.84 | *0.01 |
| Extinction | Active Lever | F | 1,13 | 0.21 | 0.66 | F | 6,78 | 38.91 | <0.0001 | F | 6,78 | 1.91 | 0.89 |
|  | Inactive Lever | F | 1,13 | 0.22 | 0.65 | F | 6,78 | 8.23 | <0.0001 | F | 6,78 | 0.81 | 0.57 |
| Memory Retrieval | Active Lever | t | 13 | 0.20 | 0.67 |  |  |  |  |  |  |  |  |
|  | Inactive Lever | t | 13 | 3.80 | 0.07 |  |  |  |  |  |  |  |  |

\*VEH > AM251 on day 8

**Figure 2-1.**
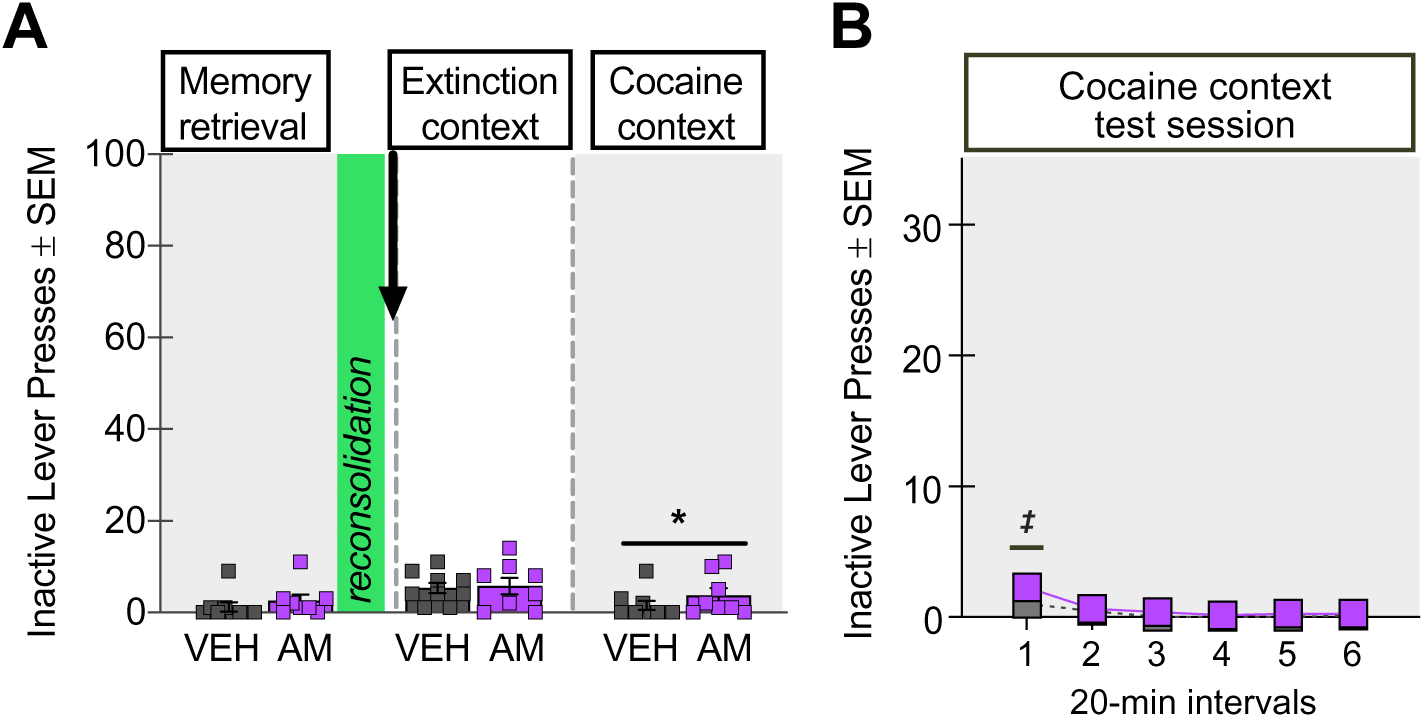
Inactive-lever responding for Experiment 2. **(A)** Inactive-lever responding was not different between the groups during the 15-min memory retrieval session (*left panel*) before administration of AM251 (AM, *n* = 8) or VEH (*n* = 9) (t_(15)_ = 0.89, p = 0.39) (*arrow*). After treatment, VEH-treated rats exhibited less inactive-lever responding in the cocaine-paired context relative to the extinction context (ANOVA, **\***context main effect, F_(1,15)_ = 6.99, p = 0.02). **(B)** Inactive-lever responding declined over time during the test in the cocaine-paired context independent of treatment (ANOVA, time main effect, F_(5,75)_ = 7.35, p < 0.0001; ***^‡^***time simple main effects, Tukey’s tests, interval 1 > 2-6, ps < 0.05).

**Table 2-1.**
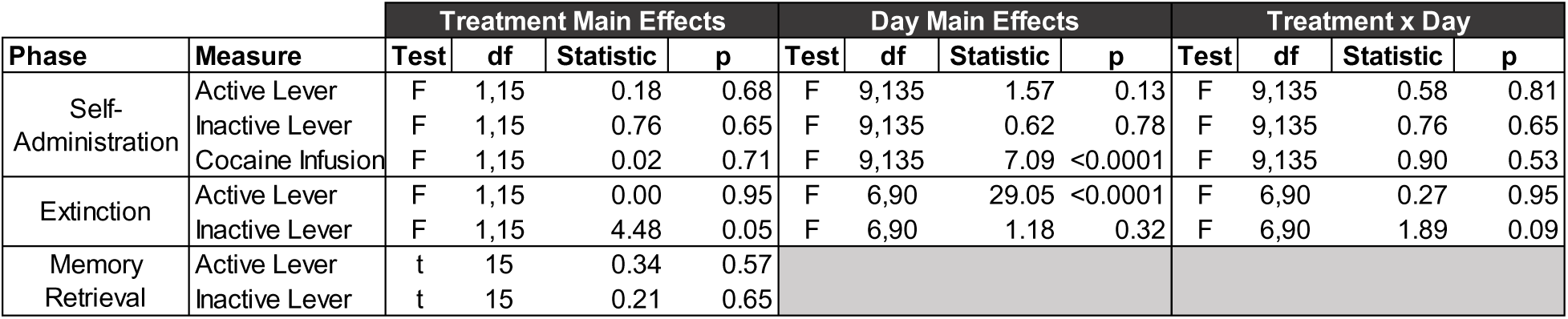
Experiment 2 Behavioral History

| Phase | Measure | Treatment Main Effects |  |  |  | Day Main Effects |  |  |  | Treatment x Day |  |  |  |
| --- | --- | --- | --- | --- | --- | --- | --- | --- | --- | --- | --- | --- | --- |
|  |  | Test | df | Statistic | p | Test | df | Statistic | p | Test | df | Statistic | p |
| Self-Administration | Active Lever | F | 1,15 | 0.18 | 0.68 | F | 9,135 | 1.57 | 0.13 | F | 9,135 | 0.58 | 0.81 |
|  | Inactive Lever | F | 1,15 | 0.76 | 0.65 | F | 9,135 | 0.62 | 0.78 | F | 9,135 | 0.76 | 0.65 |
|  | Cocaine Infusion | F | 1,15 | 0.02 | 0.71 | F | 9,135 | 7.09 | <0.0001 | F | 9,135 | 0.90 | 0.53 |
| Extinction | Active Lever | F | 1,15 | 0.00 | 0.95 | F | 6,90 | 29.05 | <0.0001 | F | 6,90 | 0.27 | 0.95 |
|  | Inactive Lever | F | 1,15 | 4.48 | 0.05 | F | 6,90 | 1.18 | 0.32 | F | 6,90 | 1.89 | 0.09 |
| Memory Retrieval | Active Lever | t | 15 | 0.34 | 0.57 |  |  |  |  |  |  |  |  |
|  | Inactive Lever | t | 15 | 0.21 | 0.65 |  |  |  |  |  |  |  |  |

**Figure 3-1.**
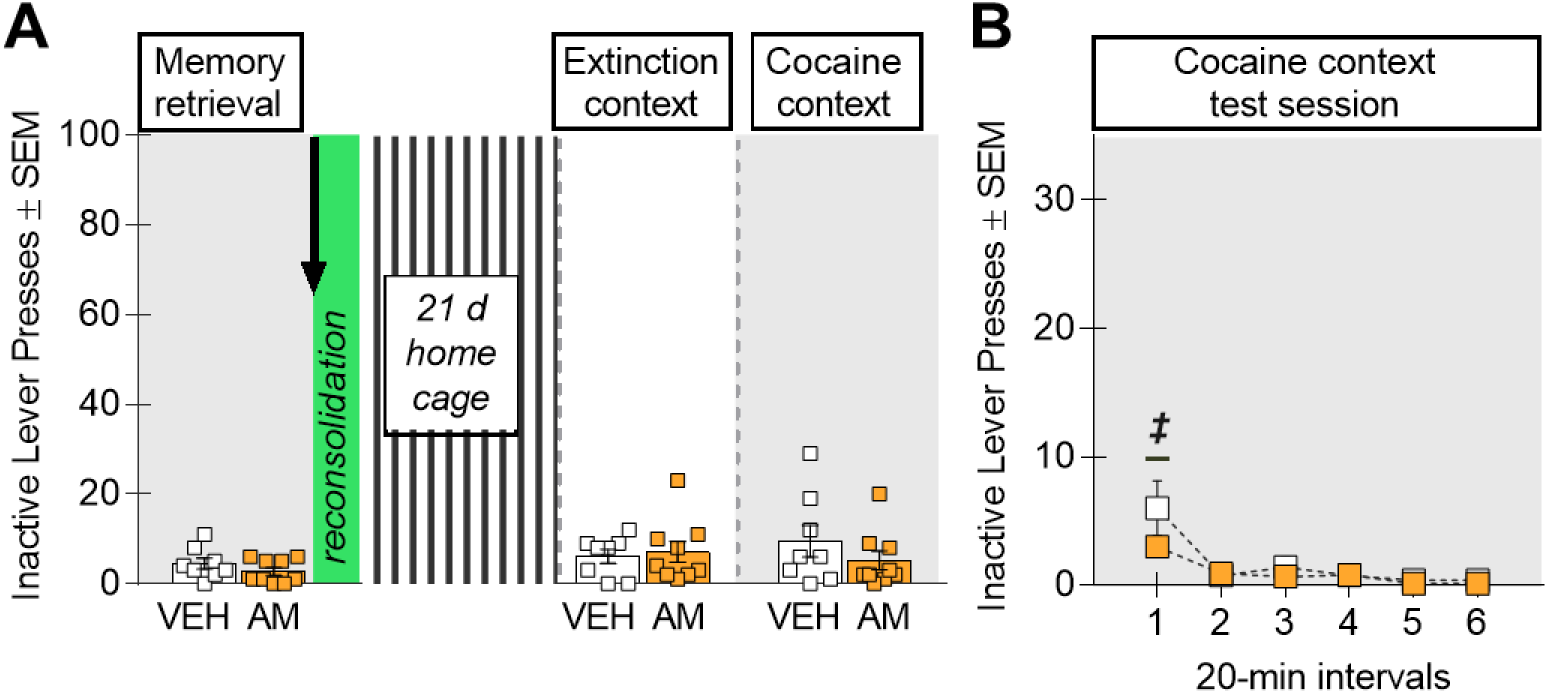
Inactive-lever responding in Experiment 3. **(A)** Inactive-lever responding was not different between the groups during the 15-min memory retrieval session (*left panel*) before administration of AM251 (AM, *n* = 9) or VEH (*n* = 8) (t_(15)_ = 1.15, p = 0.27) (*arrow*). After treatment, inactive-lever responding remained low in the extinction and cocaine-paired contexts independent of treatment (ANOVA, all Fs_(1,15)_ ≤ 1.45, p ≥ 0.25). **(B)** Inactive-lever responding declined over time in the cocaine-paired context at test independent of treatment (ANOVA, time main effect, F_(5,75)_ = 13.31, p < 0.0001; ***^‡^*** time simple main effects, Tukey’s tests, interval 1 > 2-6, ps < 0.05).

**Table 3-1.**
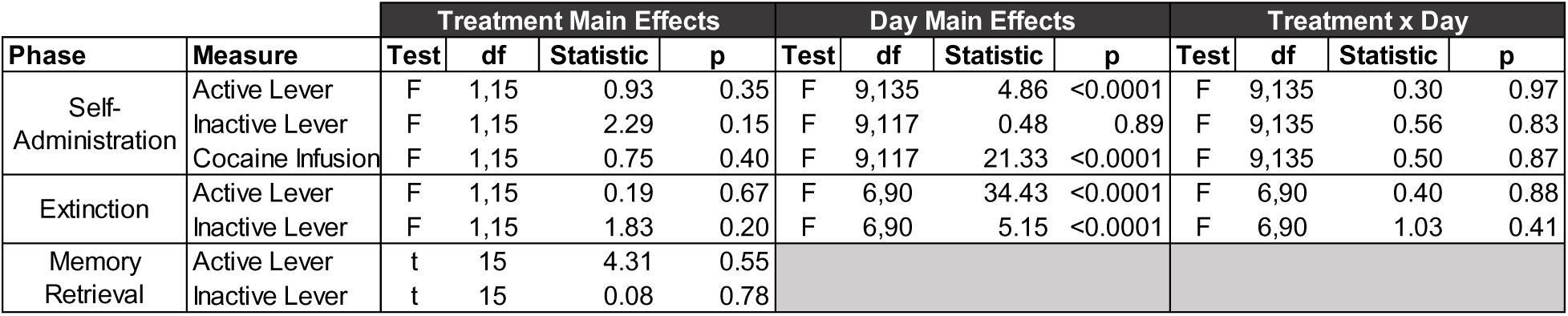
Experiment 3 Behavioral History

**Figure 4-1.**
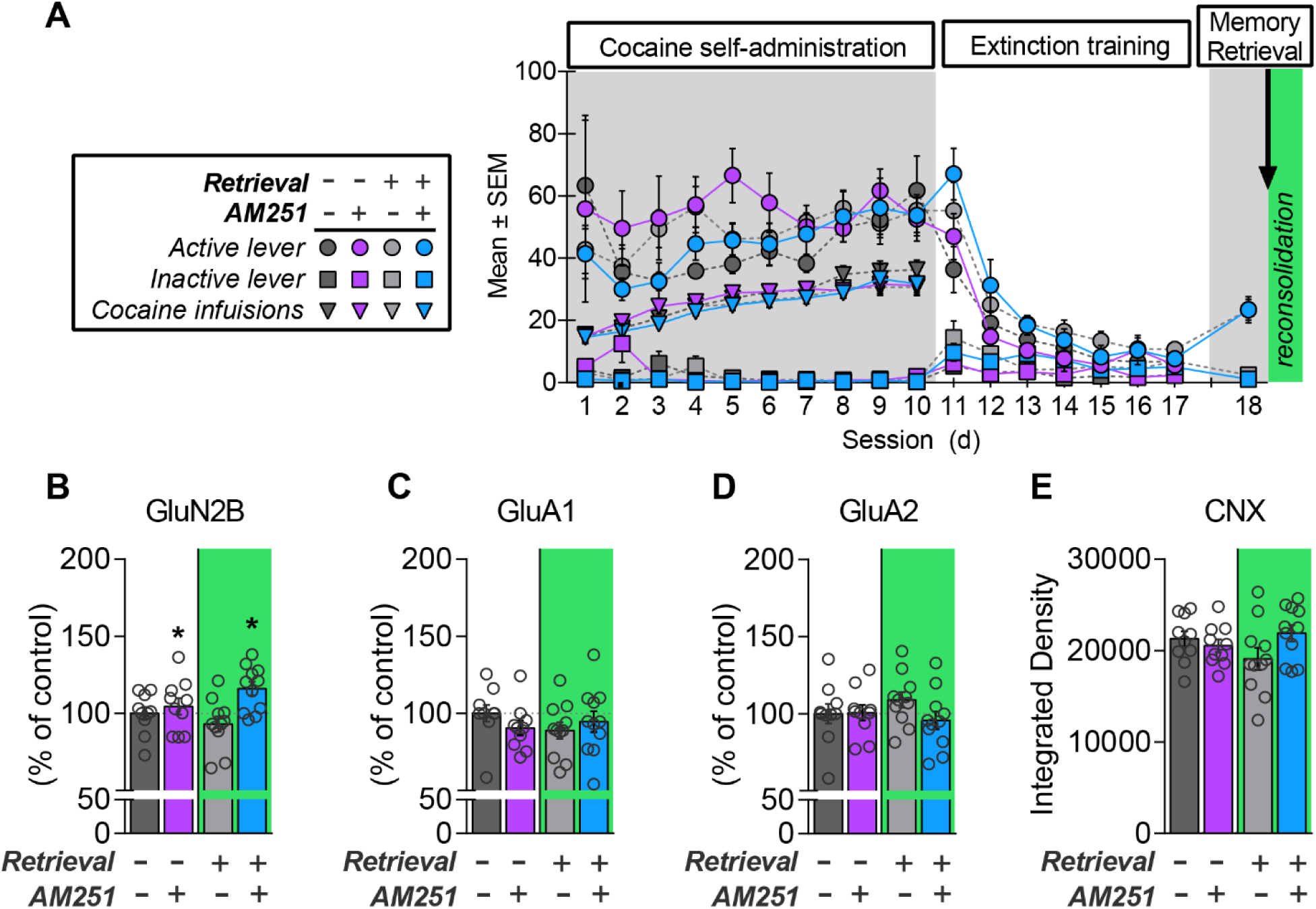
Systemic AM251 treatment increases total GluN2B, but not total GluA1 or GluA2 levels in the BLA during memory reconsolidation. **(A)** Lever responses and cocaine infusions during drug self-administration, extinction training, and the 15-min memory-retrieval session in the cocaine-paired context prior to treatment (*arrow*) in Experiment 4a. **(B)** AM251 increased total GluN2B protein levels independent of memory retrieval (ANOVA, *treatment main effect; p = 0.001). There was no effect of memory retrieval or AM251 treatment on **(C)** total GluA1 or **(D)** total GluA2 protein levels, or on **(E)** the raw integrated density values for the house-keeping protein, calnexin (CNX).

**Table 4-1.** Experiment 4 Behavioral History

| Phase | Measure | Treatment Main Effects |  |  |  | Day Main Effects |  |  |  | Treatment x Day |  |  |  |
| --- | --- | --- | --- | --- | --- | --- | --- | --- | --- | --- | --- | --- | --- |
|  |  | Test | df | Statistic | p | Test | df | Statistic | p | Test | df | Statistic | p |
| Self-Administration | Active Lever | F | 3,38 | 1.19 | 0.33 | F | 9,342 | 1.96 | 0.43 | F | 27,342 | 0.44 | 0.79 |
|  | Inactive Lever | F | 3,38 | 1.97 | 0.14 | F | 9,342 | 3.50 | <0.0001 | F | 27,342 | 2.47 | <0.0001 |
|  | Cocaine Infusion | F | 3,38 | 0.65 | 0.59 | F | 9,342 | 51.31 | <0.0001 | F | 27,342 | 0.77 | 0.79 |
| Extinction | Active Lever | F | 3,38 | 2.98 | 0.04 | F | 6,228 | 64.27 | <0.0001 | F | 18,228 | 1.56 | 0.07 |
|  | Inactive Lever | F | 3,38 | 1.49 | 0.23 | F | 6,228 | 5.47 | <0.0001 | F | 18,228 | 1.21 | 0.26 |
| Memory Retrieval | Active Lever | t | 20 | 0.94 | 0.34 |  |  |  |  |  |  |  |  |
|  | Inactive Lever | t | 20 | 1.31 | 0.27 |  |  |  |  |  |  |  |  |

**Table 5-1.**
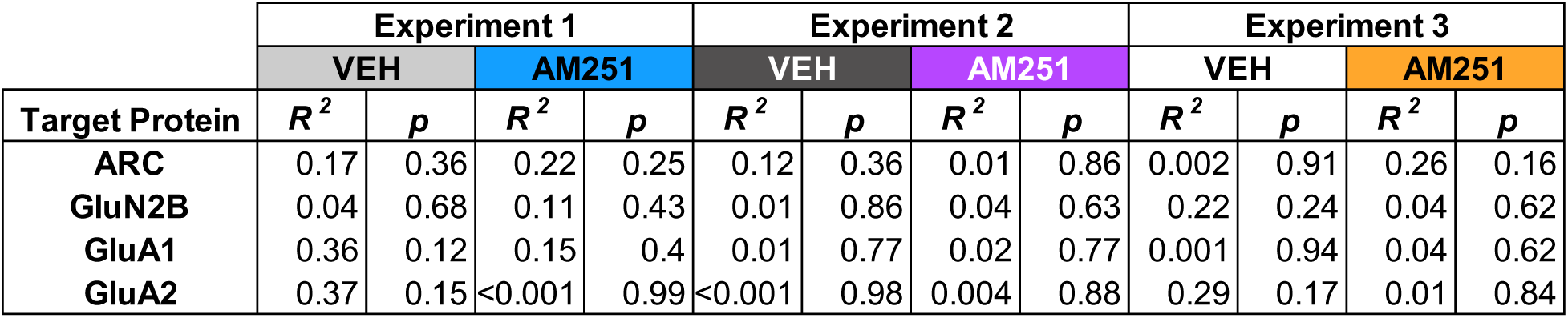
Relationship between total protein levels and active-lever presses at test

| Target Protein | Experiment 1 |  |  |  | Experiment 2 |  |  |  | Experiment 3 |  |  |  |
| --- | --- | --- | --- | --- | --- | --- | --- | --- | --- | --- | --- | --- |
|  | VEH |  | AM251 |  | VEH |  | AM251 |  | VEH |  | AM251 |  |
| | $R^2$ | $p$ | $R^2$ | $p$ | $R^2$ | $p$ | $R^2$ | $p$ | $R^2$ | $p$ | $R^2$ | $p$ |
| **ARC** | 0.17 | 0.36 | 0.22 | 0.25 | 0.12 | 0.36 | 0.01 | 0.86 | 0.002 | 0.91 | 0.26 | 0.16 |
| **GluN2B** | 0.04 | 0.68 | 0.11 | 0.43 | 0.01 | 0.86 | 0.04 | 0.63 | 0.22 | 0.24 | 0.04 | 0.62 |
| **GluA1** | 0.36 | 0.12 | 0.15 | 0.4 | 0.01 | 0.77 | 0.02 | 0.77 | 0.001 | 0.94 | 0.04 | 0.62 |
| **GluA2** | 0.37 | 0.15 | <0.001 | 0.99 | <0.001 | 0.98 | 0.004 | 0.88 | 0.29 | 0.17 | 0.01 | 0.84 |

**Figure 6-1.**
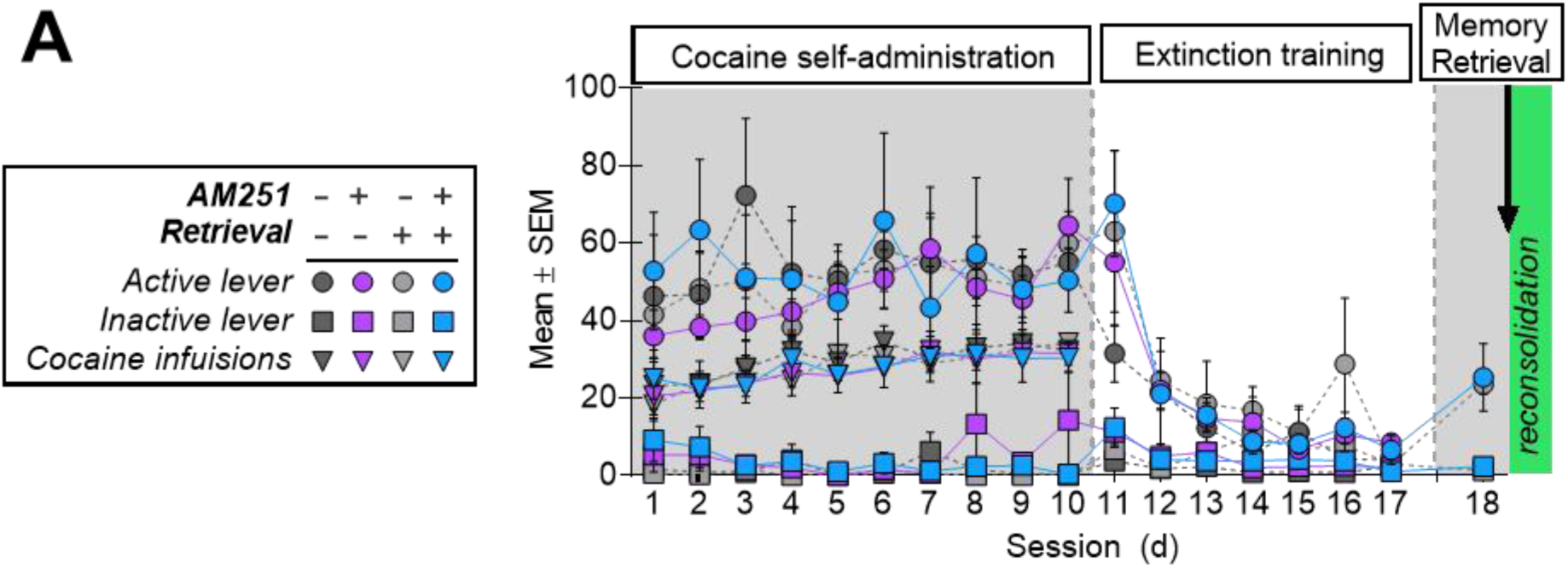
Behavioral history in Experiment 5. **(A)** Lever responses and cocaine infusions during drug self-administration, extinction training, and the 15-min memory-retrieval session in the cocaine-paired context prior to treatment (*arrow*). Statistics are provided in Table 6-1.

**Table 6-1.** Experiment 5 Behavioral History

| Phase | Measure | Treatment Main Effects |  |  |  | Day Main Effects |  |  |  | Treatment x Day |  |  |  |
| --- | --- | --- | --- | --- | --- | --- | --- | --- | --- | --- | --- | --- | --- |
|  |  | Test | df | Statistic | p | Test | df | Statistic | p | Test | df | Statistic | p |
| Self-Administration | Active Lever | F | 3,13 | 0.29 | 0.83 | F | 9,117 | 0.75 | 0.66 | F | 27,117 | 0.69 | 0.86 |
|  | Inactive Lever | F | 3,13 | 1.13 | 0.37 | F | 9,117 | 0.77 | 0.65 | F | 27,117 | 1.19 | 0.26 |
|  | Cocaine Infusion | F | 3,13 | 0.11 | 0.95 | F | 9,117 | 7.32 | <0.0001 | F | 27,117 | 0.55 | 0.96 |
| Extinction | Active Lever | F | 3,12 | 0.10 | 0.96 | F | 6,72 | 22.61 | <0.0001 | F | 6,72 | 0.21 | 1.00 |
|  | Inactive Lever | F | 3,12 | 1.04 | 0.41 | F | 6,72 | 6.63 | <0.0001 | F | 6,72 | 1.06 | 0.41 |
| Memory Retrieval | Active Lever | t | 7 | 5.00 | 0.06 |  |  |  |  |  |  |  |  |
|  | Inactive Lever | t | 7 | 3.42 | 0.11 |  |  |  |  |  |  |  |  |

**Table 6-2.** Outward current frequency and amplitude with aCSF

| Recording Condition | Measure | Treatment Main Effects |  |  |  | Day Main Effects |  |  |  | Treatment x Day |  |  |  | Post-hoc effects |
| --- | --- | --- | --- | --- | --- | --- | --- | --- | --- | --- | --- | --- | --- | --- |
|  |  | Test | df | Statistic | p | Test | df | Statistic | p | Test | df | Statistic | p | Sidak's or Dunn's test, p < 0.05 |
| aCSF | sEPSC Frequency (+30 mV) | F | 1,20 | 0.07 | 0.80 | F | 1,20 | 7.54 | 0.01 | F | 1,20 | 3.01 | 0.10 | NA |
|  | Frequency Distribution |  |  |  |  |  |  |  |  | H | 3 | 259.70 | < 0.0001 | Memory retrieval + AM251 > all other groups |
|  | sEPSC Amplitude (+30 mV) | F | 1,20 | 0.02 | 0.89 | F | 1,20 | 0.16 | 0.70 | F | 1,20 | 1.43 | 0.25 | NA |
|  | Amplitude Distribution |  |  |  |  |  |  |  |  | H | 3 | 16.36 | 0.00 | No retrieval + AM251 < all other groups |

**Table 7-1.** Outward current frequency and amplitude with AP5

| Recording Condition | Measure | Treatment Main Effects |  |  |  | Day Main Effects |  |  |  | Treatment x Day |  |  |  | Post-hoc effects<br>Sidak's or Dunn's test,<br>p < 0.05 |
| --- | --- | --- | --- | --- | --- | --- | --- | --- | --- | --- | --- | --- | --- | --- |
|  |  | Test | df | Statistic | p | Test | df | Statistic | p | Test | df | Statistic | p |  |
| AP5 | sEPSC Frequency (+30 mV) | F | 1,43 | 2.67 | 0.11 | F | 1,43 | 21.15 | <0.0001 | F | 1,43 | 10.14 | 0.00 | Memory retrieval + AM251 > all other groups |
|  | Frequency Distribution (+30 mV) |  |  |  |  |  |  |  |  | H | 3 | 180.70 | < 0.0001 | Memory retrieval + AM251 > all other groups |
|  | sEPSC Amplitude (+30 mV) | F | 1,44 | 0.41 | 0.53 | F | 1,44 | 0.25 | 0.62 | F | 1,43 | 1.67 | 0.20 | NA |
|  | Amplitude Distribution (+30 mV) |  |  |  |  |  |  |  |  | H | 3 | 0.46 | 0.93 | NA |

